# Micropatterned neural induction with heat-inactivated extracellular matrix protein by on-demand high-speed laser

**DOI:** 10.1101/2025.07.08.663334

**Authors:** Yohei Hayashi, Junichi Matsumoto, Kimio Sumaru

## Abstract

Microenvironmental heterogeneity in cultured cells can compromise cell quality, reduce experimental reproducibility, and weaken the confidence of cell therapeutic efficacy. Although micropatterned cell cultures are more homogeneous, conventional micropatterning methods lack flexibility. In this study, we develop a micropatterning technology by denaturing extracellular matrix (ECM) proteins in specific areas through heat inactivation using a high-speed laser via a light-responsive polymer layer. We have successfully seeded human induced pluripotent stem cells (hiPSCs) in flexible patterns and examine their neural induction in circular geometries of varying diameters. Size-dependent and cell-autonomous neural structures are formed on this substrate when hiPSCs differentiate into neural lineages in circles of different diameters. This self-organized pattern results from the mitotic orientation and localization of differentiating cells. Furthermore, teratogenic substances can modulate these patterns. Laser-induced heat inactivation of the ECM on culture substrates enables on-demand patterning, facilitating studies of cell-autonomous tissue formation, the effect of teratogenic substances, and precise tissue engineering in regenerative medicine.

**Table of Content:** A lithography-, hydrogel-, and PDMS-free micropatterning technology is developed to enable flexible and rapid cell patterning directly onto culture substrates via high-speed laser irradiation. Using this technology, size-dependent, cell-autonomous tissue formation from human induced pluripotent stem cells were driven by mitotic orientation and spatial localization of differentiating cells. These self-organized structures can be applied for safety tests of teratogenic substances.

**ToC figure:** 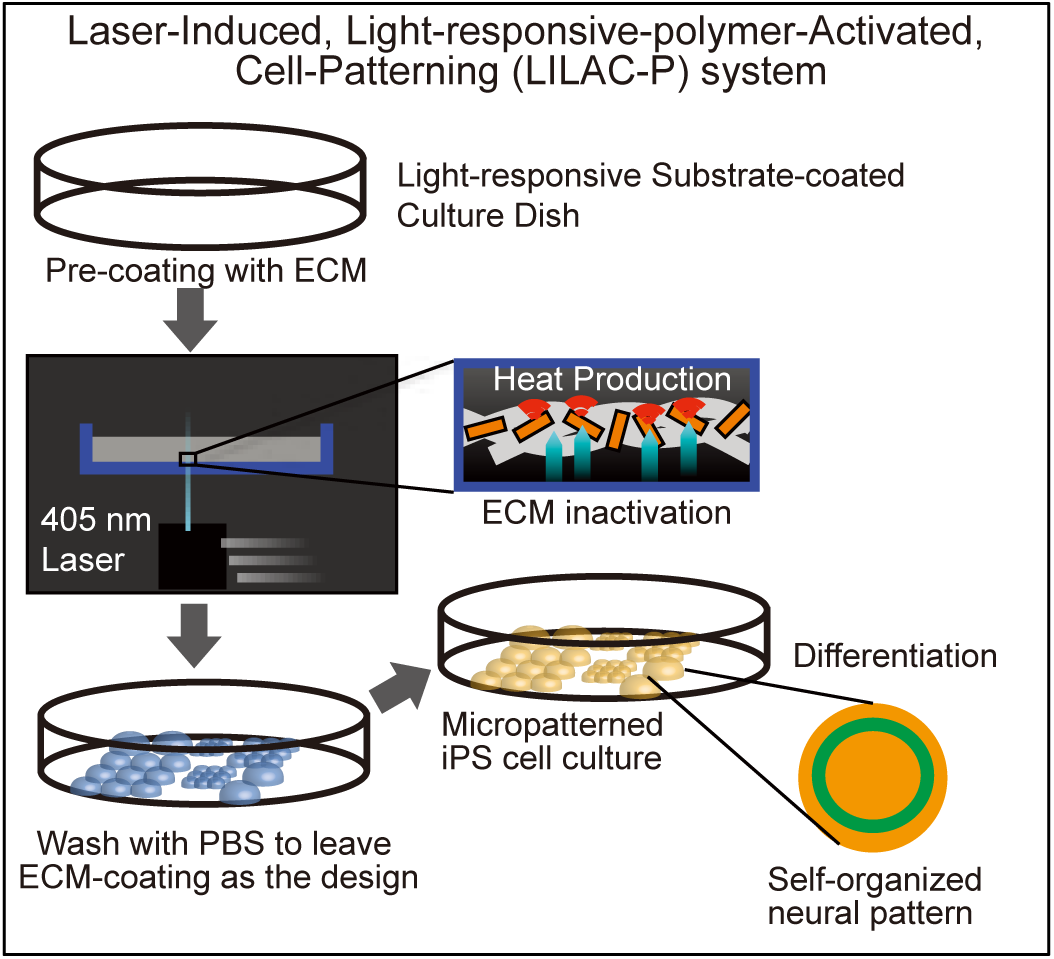

## 1. Introduction

The cellular microenvironment sustains the formation of tissues and organs and regulates cell status.^[1–4]^ Cell cultures can mimic such microenvironments, allowing a detailed examination of cellular responses to diverse stimuli.^[5–7]^ However, controlling the microenvironment of individual cells using conventional culture methods is challenging, leading to heterogeneity and ambiguity in their cellular state that cause high variability and low reproducibility of experimental results in basic research and drug discovery. Furthermore, in regenerative medicine, such microenvironmental uncertainty can reduce the therapeutic efficacy of cellular products.^[8–9]^

Micropatterned culture methods efficiently control the cellular microenvironment, facilitating *in vitro* analyses of cellular state *in vitro* and the induction of novel cell types and tissues under these culture conditions.^[10–15]^ Various micropatterned culture methods have been developed,^[16–19]^ with the main techniques including: 1) photolithography, in which the surface of a material coated with a photosensitive material is exposed in a specific pattern, producing exposed and unexposed areas;^[20–22]^ 2) stamping soft lithography, in which a hydrogel or other surface is molded, modified, and microfabricated to produce a pattern of exposed and unexposed areas;^[23–25]^ 3) microfluidics, in which structures with width and depth ranging from a few to several hundred microns are fabricated through semiconductor, precision machining, or optical fabrication technologies.^[26–27]^ Although these methods are widely used, micropatterning fabrication requires specialized skills and may lack flexibility, such as the ability to customize new patterns. Additionally, the culture scale of these micropatterning techniques is typically too small to control the microenvironment particularly for regenerative medicine, where high cell yields are required. There is a growing demand for technology that enables biologists and cell culture technicians, regardless of manufacturing expertise, to easily and flexibly perform micropatterning. Additionally, such a system should allow scalability for large-scale cell culture.

Therefore, we aimed to develop a micropatterned cell culture method that differs from conventional technologies. In our previous study, a Laser-Induced, Light-responsive polymer-Activated Cell Killing (LILACK) system was developed to enable high-speed and on-demand adherent cell sectioning and purification.^[28]^ This LILACK system employs a 405 nm visible laser beam to produce localized heating at the specific irradiated area of a light-responsive thin layer composed of poly[(methyl methacrylate)-co-(Disperse Yellow 7 methacrylate)]. The laser does not directly kill cells, but the energy of the irradiated laser is efficiently converted to heat through the trans-cis-trans photoisomerization of azobenzene moieties without photolysis of the polymer.^[29]^ Laser irradiation at a speed exceeding 100 mm s^−1^ and with a beam diameter of approximately 10 µm accurately produced localized heating above 100 °C on the surface of the light-responsive-polymer-coated dishes. This polymer is free from fluorescence emission and absorbance across most of the visible spectrum, thereby allowing unobstructed cell observations. This LILACK system enables *in situ* selection of adherent cells within an acceptable timescale by precisely and rapidly scanning a well-focused visible laser through a light-responsive polymer layer, without inducing cytotoxicity, enabling automatic, label-free cell purification combined with efficient imaging analysis based on deep machine learning methods. This technology has been demonstrated for the label-free purification of undifferentiated hiPSCs by depleting spontaneously-differentiated cells in culture. ^[28]^ Recently, another research group utilizes this technology for the label-free purification of hiPSC-derived cardiomyocytes from other heterogenous cell types. ^[30]^

In the present study, we modified LILACK to irradiate culture substrates coated with various extracellular matrix (ECM) proteins to denature the pre-coated ECM in specific irradiated areas. When cultured cells were subsequently seeded onto the surface, adhesion occurred only in areas that had not been irradiated by the laser. Based on this principles, we established a Laser-Induced, Light-responsive-polymer-Activated, Cell-Patterning (LILAC-P) system that enables high-speed, on-demand patterning of the entire culture dish by automatic laser irradiation guided by image data. This technology enables cell patterning and the analysis of changes caused by the pattern shape in cultured human induced pluripotent stem cells (hiPSCs) and their differentiated progeny.

## 2. Results

### 2.1. Thermal denaturation of ECM proteins and patterning of cell seeding by high-speed laser irradiation mediated with light-responsive polymers

We initially assessed the feasibility of micropatterned cell seeding via heat-induced denaturation of the ECM proteins on culture dishes coated with light-responsive polymers, using high-speed laser irradiation (**Figure 1A**). In this scheme, the entire culture dish was pre-coated with ECM proteins dissolved in Dulbecco’s phosphate-buffered saline (D-PBS); after a wash with D-PBS, the entire culture dish was laser-irradiated based on input from a black and white image file (**Figure 1B**). The dish was then washed with PBS, and hiPSCs were seeded. First, the intensity and speed of the laser were examined to optimize the irradiation conditions. An effective cell seeding pattern was observed at laser output power ≥0.8 W (**Figure 1C**) and laser speeds of ≤120 mm s^−1^ **Figure 1D**). Next, using these optimized laser irradiation conditions, we demonstrated various patterning of the entire 35 mm dish with seeded hiPSCs. Depending on the dish design, the 35 mm dishes were filled with hiPSCs in various patterns: 812 (28 × 29) pieces of 300 µm squares on one side (**Figure 1E**), a mesh pattern covering the entire bottom surface (**Figure 1F**), string shapes of 60 µm width and length up to 1 cm (**Figure 1G**), or character-shaped patterns (**Figure 1H**). These results demonstrate that cell seeding patterning can be customized using a heat-denatured polymer substrate and high-speed laser irradiation (LILAC-P).

**Figure 1.**
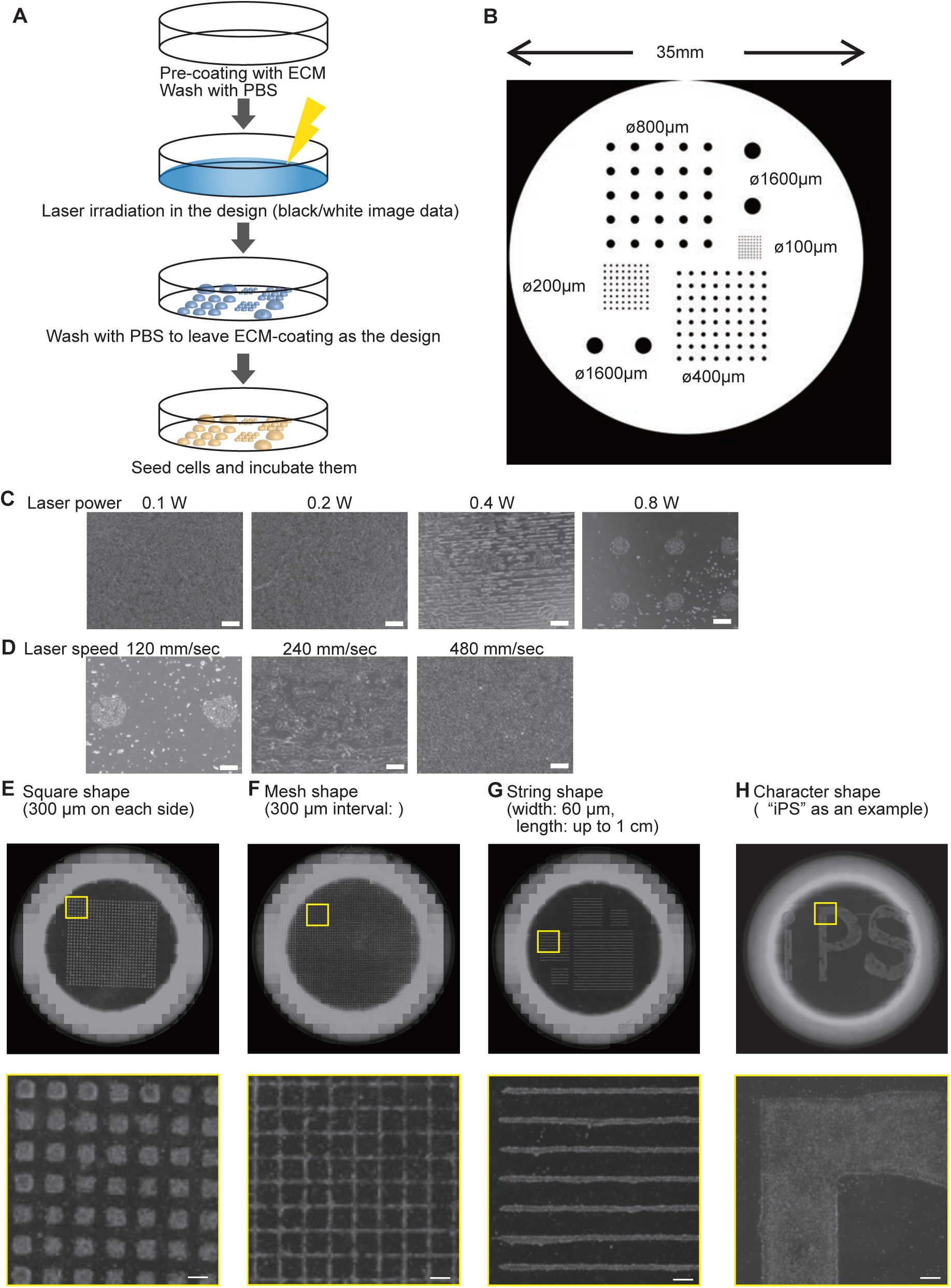
Proof of concept of Laser-Induced, Light-responsive-polymer-Activated, Cell-Patterning (LILAC-P). (A) Schematics of the laser irradiation and the cell seeding patterning. (B) Representative laser irradiation pattern design. Different size of black dots (100, 200, 400, 800, and 1600 µm diameters) were placed in a 35 mm circle. The laser machine irradiates only the white area. (C) LILAC-P with different laser powers of 0.1, 0.2, 0.4, and 0.8 W. Scale bars, 400 µm. (D) LILAC-P with different laser speeds of 120, 240, and 480 mm s^−1^. Scale bars, 400 µm. (E) Representative example of micropattern of a whole dish (upper) and a magnified view of the yellow square region (bottom),each measuring 300 µm per side. Scale bar, 300 µm. (F) Representative example of micropattern of a whole dish (upper) and a magnified view of the yellow square region (bottom) of mesh, with 300 µm spacing. Scale bar, 300 µm. (G) Representative example of micropattern of a whole dish (upper) and a magnified view of the yellow square region (bottom) of strings, 60 µm wide and length up to 1 cm. Scale bar, 300 µm. (H) Representative example of micropattern of a whole dish (upper) and a magnified image of the yellow square region (bottom) of characters (“iPS”). Scale bar, 300 µm.

Next, we examined whether this LILAC-P technology was mediated by the thermal denaturation of ECM proteins on a culture dish coated with a light-responsive polymer by high-speed laser irradiation. To begin the experiment, we examined how the binding ability of laminin to integrin changed following laser irradiation using fluorescently labeled purified recombinant integrin receptors in solid-phase assays (**Figure S1A**).^[31]^ Consistent with the earlier cell seeding study, clear integrin-binding patterning was observed at a laser speed of 120 mm s^−1^, but failed under higher-speed conditions (**Figure S1B**). The fluorescence of the area irradiated at a laser speed of 120 mm s^−1^ was similar to the non-fluorescently labeled recombinant integrin (**Figure S1C**). These results indicated that sufficient laser irradiation completely eliminated the integrin-binding capacity of the coated laminin. Further examination of the laser power revealed similar clear integrin-binding patterning at 0.8 W (**Figure S1D**). The fluorescence of the non-laser- and laser-irradiated areas revealed a marked difference in fluorescence intensity (**Figure S1E**), indicating that this laser irradiation causes a loss of integrin-binding ability within the targeted area.

We further applied biotinylated laminin under identical conditions, followed by treatment with streptavidin-labeled allophycocyanin (APC) to identify undenatured laminin, based on the strong and specific binding affinity between biotin and streptavidin (**Figure S2A**). The laser-irradiated dishes showed clear fluorescent patterning of APC corresponding to the site of laser irradiation at 0.8 W and 120 mm s^−1^ (**Figure S2B**). APC fluorescence was consistently lower in laser-irradiated dishes compared to the non-laser irradiated dishes across the board (**Figure S2C**). Thus, laser irradiation results in the loss of the streptavidin-binding ability of biotin, which modifies laminin. Overall, these findings indicate that high-speed laser irradiation induces heat denaturation of ECM proteins on culture dishes coated with light-responsive polymers, leading to the loss of integrins and, consequently, cell-binding ability.

### 2.2. Behavior of hiPSCs on cell-seeded patterns formed via thermal denaturation of ECM proteins using high-speed laser irradiation

We evaluated whether hiPSCs maintained normal behavior on these LILAC-P-generated micropatterns when seeded in circular patterns with diameters of 100, 200, 400, 800, and 1600 µm. After culturing cells for 2 days, we examined the expression of OCT3/4 (**Figure S3A**), NANOG (**Figure S3B**), TRA-1-60 cell surface marker (**Figure S3C**), and SSEA4 cell surface marker (**Figure S3D**), which are markers of undifferentiated hiPSCs.^[32]^ Micropatterned seeded hiPSCs were positive for all the markers, indicating that their self-renewal was maintained on patterns generated by the LILAC-P system.

Next, we examined whether hiPSCs could differentiate into specific cellular lineages based on the LILAC-P-generated micropatterns. HiPSCs were seeded onto these patterns and cultured overnight. The next day, these cells were induced to differentiate into neuroectodermal (Figure 2A), mesodermal (**Figure S4A**), or endodermal cells (**Figure S5A**) using a specific culture medium.^[33]^ Seven days after seeding, the cells were analyzed by immunostaining. When hiPSCs were differentiated into the neuroectoderm using the dual SMAD inhibition method, the integrity of the circular pattern was generally preserved across all diameters (Figure 2B).^[34]^ However, the 100 µm diameter circular pattern was relatively distorted, likely attributed to the limited space available in relation to the cell size and laser irradiation width, and was consequently excluded from future analysis. Under these differentiation conditions in patterned cultures, OCT3/4 expression was lost, and highly efficient expression of PAX6, a marker of the neuroectoderm, was observed, similar to that in monolayer control conditions. Thus, hiPSCs in the LILAC-P system can efficiently differentiate into neuroectoderm. During the process of inducing mesoderm, some OCT3/4-negative, HAND1-positive cardiovascular mesoderm cells appeared across all pattern sizes (**Figure S4B**). However, these cells deviated from the seeding pattern owing to migration during the culture process. During endoderm induction, some OCT3/4-negative and SOX17-positive endodermal cells appeared across all pattern sizes (**Figure S5B**). As in the mesodermal differentiation process, these cells deviated from the seeding pattern because of migration during the culture process. In summary, hiPSCs can differentiate into specific cellular lineages in response to medium stimuli using the LILAC-P system. In contrast, the culture pattern can be disrupted in cells with a high migratory ability. Therefore, in this study, we focused on the induction of differentiation into neural lineages, in which the seeding pattern was maintained during long-term culture.

**Figure 2.**
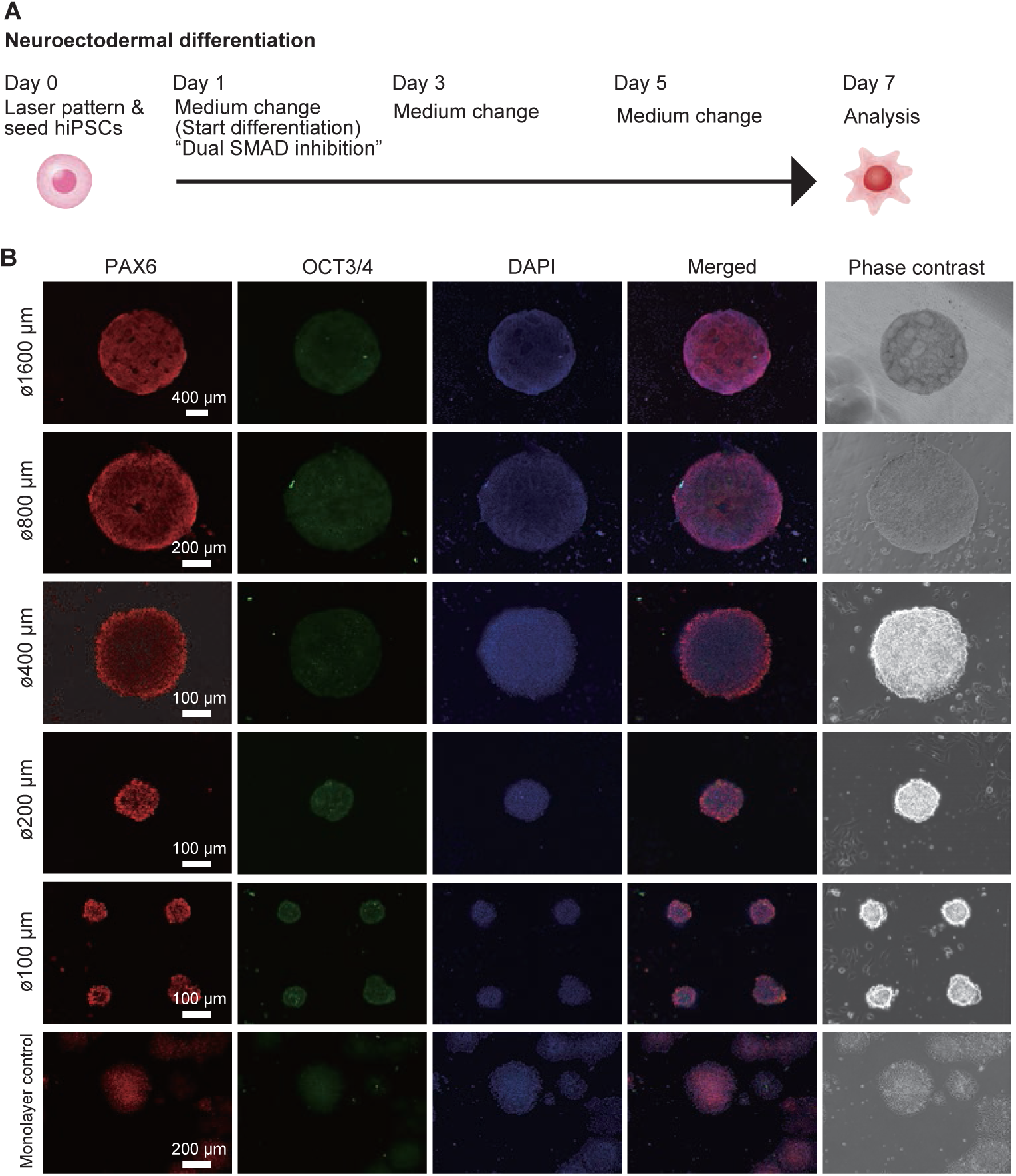
Neuroectoderm differentiation of hiPSCs cultured on dishes using the LILAC-P system. (A) Schematics of neuroectodermal differentiation in culture dishes with LILAC-P system for one week. (B) Representative images of PAX6 (red), OCT3/4 (green), and DAPI (blue) detected with immunocytochemistry. Merged and phase contrast images from the right at different diameters (1600, 800, 400, 200, and 100 µm) alongside monolayer control samples. The size of each scale bar is shown in the figure.

### 2.3. Self-organized cortical layer formation of polarized neuroectoderm from hiPSCs cultured with the LILA-P system

Under conditions of neuroectodermal induction in micropatterned cultures using the LILAC-P system, the expression of other neural markers was examined. N-cadherin (CDH2), a neural cell adhesion molecule, is specifically expressed in the nervous system.^[35]^ Notably, a constant ring-shaped expression pattern of N-cadherin was obtained for each diameter pattern; however, almost no positive cells were obtained on day 7 of induction under monolayer control conditions (Figure 3A). Forkhead box G1 (FOXG1) is one of the earliest transcription factors expressed during early neurogenesis. It responds to various signaling pathways and is crucial in forebrain development.^[36–37]^ Herein, simple monolayer cultures and 1600 µm diameter cells were uniformly positive for FOXG1, whereas patterns below 800 µm showed a specific expression pattern in the outer layers (Figure 3B), contrasting with the uniform expression pattern of the co-stained PAX6 protein. Next, we changed the cell line and performed experiments using the PAX6-TEZ, which expresses tdTomato red fluorescent protein in response to PAX6 protein expression and was generated in our previous study.^[38]^ On day 7 of differentiation, similar to the PAX6 immunocytochemistry, PAX6-tdTomato positive cells were uniformly obtained for each diameter in a circular pattern. These cells were then examined for NESTIN, an intermediate-diameter filament expressed in neural progenitor cells.^[39]^ Circular patterns with diameters of 400 and 800 µm yielded NESTIN expression patterns oriented from the outside toward the inside. In contrast, in monolayer cultures and in 200 and 1600 µm diameter cells, the NESTIN filaments were randomly oriented in individual cells (Figure 3C). These results indicate that micropattern culture of hiPSCs promotes the expression of some neuronal differentiation markers, generating self-organized patterns of neuronal structures, and that the expression patterns of specific differentiation markers change according to pattern size.

**Figure 3.**
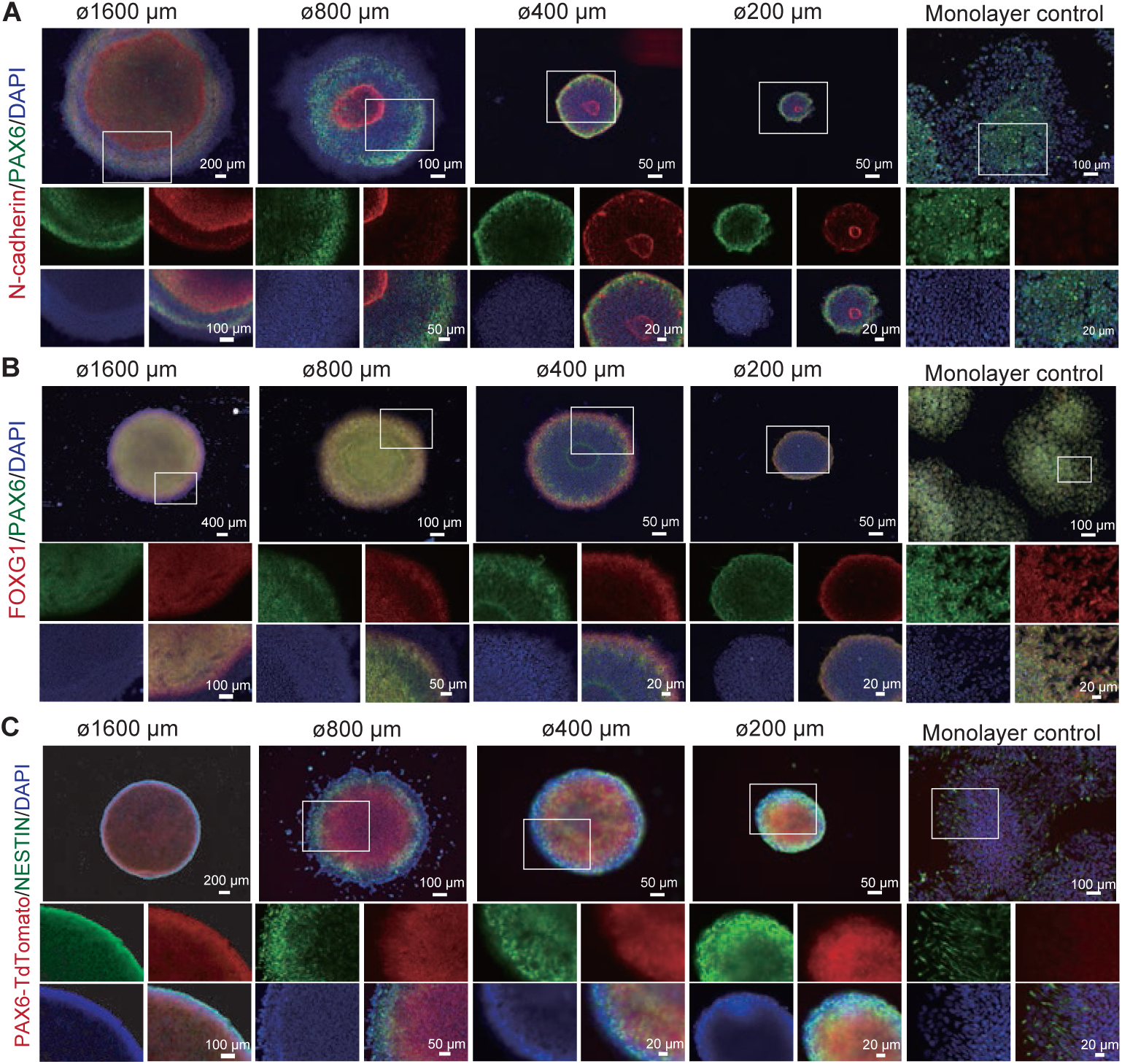
Self-organized neural induction in micropatterned culture generated with LILAC-P system. (A) Representative images of N-Cadherin (red), PAX6 (green), and DAPI (blue) at different diameters (1600, 800, 400, and 200 µm, as well as monolayer control from left to right). In the smaller panels, expression of each individual marker and a merged image of the enlargements of inserts (white square) in the upper panels are shown. (B) Representative images of FOXG1 (red), PAX6 (green), and DAPI (blue) at different sizes (1600, 800, 400, and 200 µm, as well as monolayer control from left to right). In the smaller panels, expression of each individual marker and a merged image of the enlargements of inserts (white square) in the upper panels are shown. (C) Representative images of Pax6-TEZ (red), Nestin (green), and DAPI (blue) at different diameters (1600, 800, 400, and 200 µm, as well as monolayer control from left to right). In the smaller panels, expression of each individual marker and a merged image of the enlargements of inserts (white square) in the upper panels are shown.

We then examined the neuronal differentiation patterns in long-term cultures of these seeding patterns after 1, 2, and 3 weeks of culture (Figure 4A). First, TUJ1 (TUBB3), a neuron-specific tubulin, and TBR1, a marker of post-mitotic projection neurons, were examined using immunocytochemistry (Figure 4B). Although TBR1-positive cells appeared only after 3 weeks of culture in the monolayer control conditions, their uniform expression started after 1 week of culture in the micropattern conditions. The expression of TUJ1 protein was uniformly positive after 3 weeks of culture in the monolayer control conditions, whereas a specific expression pattern in the outer layers was observed in weeks 1 to 3. Next, SOX2, a transcription factor regulating neural stem cells, and TBR2, a marker of post-mitotic projection neurons, were examined (Figure 4C). SOX2-positive cells were uniformly observed in the monolayer cultures after 1 week. After 2 and 3 weeks, the expression of SOX2 decreased in the monolayer culture; however, in the pattern culture, its expression remained restricted to the outer layer of the cells. In contrast, the TBR2 expression was unclear in these cultures. The expression of BRN2 (POU3F2) and reelin, which are crucial for the laminar brain structure and telencephalon formation,^[40]^ was examined. Reelin and BRN2 were not expressed in monolayer cultures at 1 week but became detectable after 2 weeks (Figure 4D). In the patterned cultures, BRN2 was expressed in the outermost layer from the first week, whereas in the inner layer of the 400 µm diameter culture, it was expressed in the form of another ring. After the second week, it was randomly expressed in inner cells. These results indicate that culturing on the seeding pattern promotes neuronal differentiation and generates structured layered patterns depending on the size of the micropatterns.

**Figure 4.**
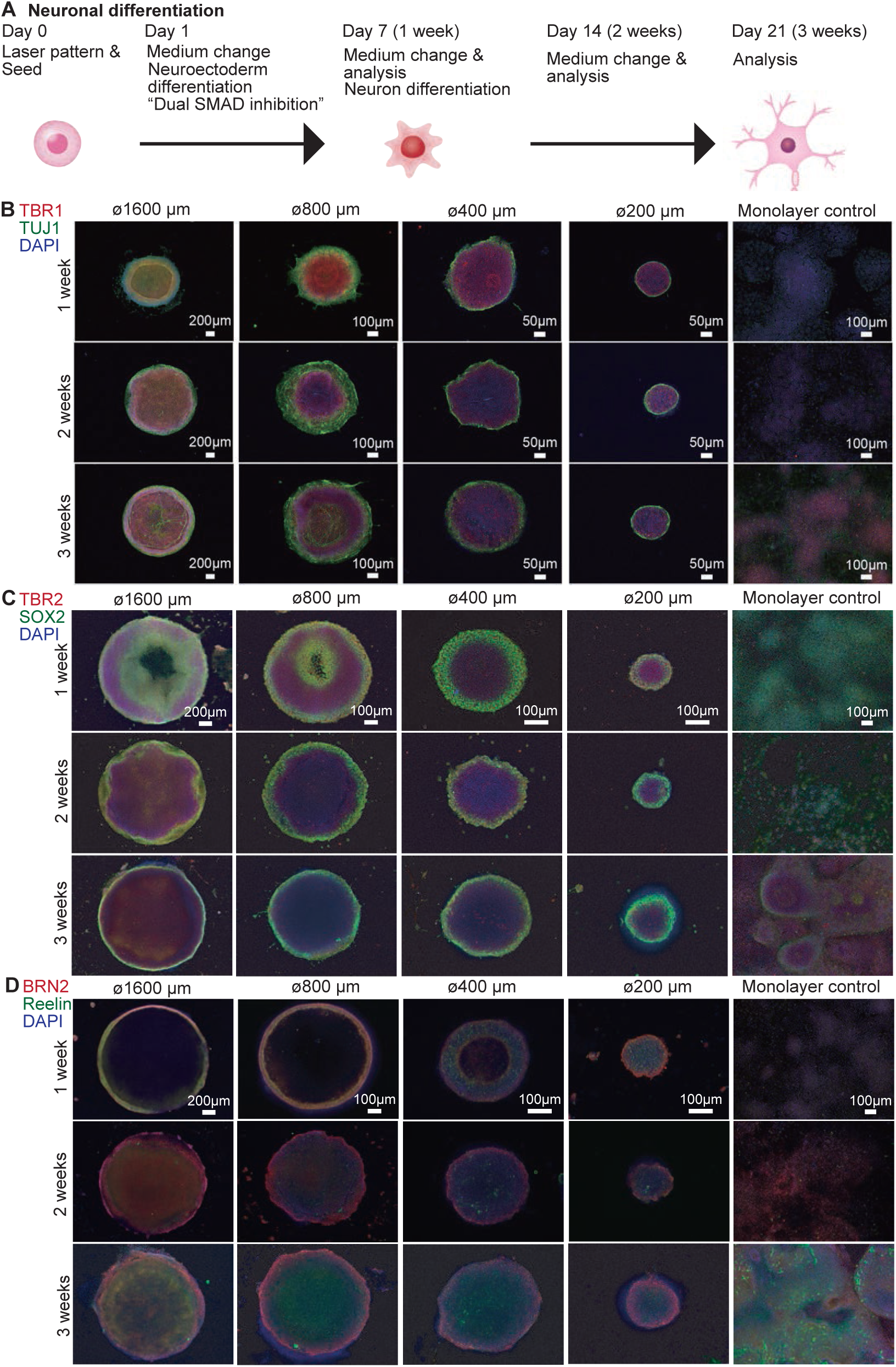
Self-organized neuronal structure in long-term micropatterned culture. (A) Schematics of neuronal differentiation in culture dishes with the LILAC-P system up to 3 weeks. (B) Representative images of TBR1 (red), TUJ1 (green), and DAPI (blue) detected with immunocytochemistry on 1-, 2-, and 3-week samples. Merged and phase contrast images from the right at different diameters of 1600, 800, 400, and 200 µm and monolayer control samples. The size of each scale bar is mentioned in each image. (C) Representative images of TBR2 (red), SOX2 (green), and DAPI (blue) detected with immunocytochemistry on 1-, 2-, and 3-week samples. Merged and phase contrast images from the right at different sizes of 1600, 800, 400, and 200 µm diameters and monolayer control samples. The size of each scale bar is mentioned in each image. (D) Representative images of BRN2 (red), Reelin (green), and DAPI (blue) detected with immunocytochemistry on 1-, 2-, and 3-week samples. Merged and phase contrast images from the right at different sizes of 1600, 800, 400, and 200 µm diameters and monolayer control samples. The size of each scale bar is mentioned in each image.

### 2.4. Autonomous layer-oriented cell division in micropattern cultures

As described above, self-organized layers of neuronal differentiation were generated in a micropattern system. To explore the mechanisms underlying the formation of these self-organized layers, we examined cell division activity during neural differentiation in the micropatterned culture. The PAX6-TEZ hiPSC line was used to ensure neural differentiation by monitoring tdTomato fluorescence and examining the expression of phospho-histone H3 (Ser10)(phospho-H3), a marker of cell division (Figure 5A). ^[39]^ In monolayer cultures, the expression of tdTomato was not evident during the first week; however, from the second week onward, it was expressed by the majority of the cells. Phospho-H3-positive cells were randomly selected under monolayer conditions. In contrast, in the micropatterned cultures, tdTomato-positive cells were uniformly maintained until the third week for all seeding patterns. The phospho-H3-positive cells were localized on the outside of the 800 and 1600 µm diameter seeding patterns during all weeks; however, positive cells formed clusters on the inside of the 200 and 400 µm diameter seeding patterns. These results indicate that cell division depends on the micropattern size during neuronal differentiation.

**Figure 5.**
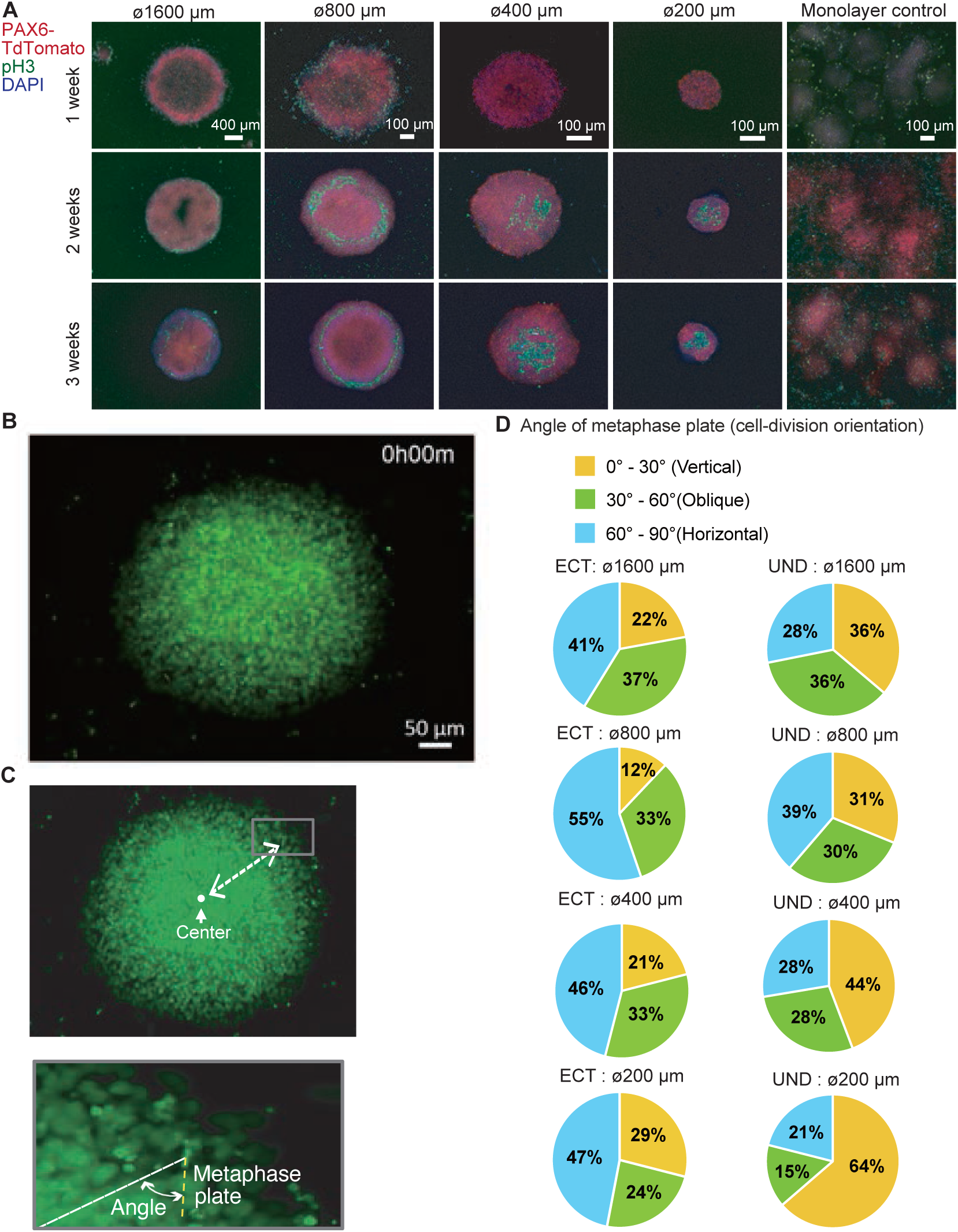
Layer-oriented cell division during ectoderm differentiation on a laser-irradiated dish. (A) Representative images of PAX6-tdTomato (red), phospho-Histone 3 (ser10) (pH3) (green), and DAPI (blue) detected with immunocytochemistry on 1-, 2-, and 3-week samples. Merged and phase contrast images from the right at different sizes of 1600, 800, 400, and 200 µm diameters and monolayer control samples. (B) Representative image frame from the time-lapse movie showing neuroectoderm differentiation in an area of 400 µm on the laser-irradiated dish. (C) Schematics illustrating the method to measure the cell-division angle during metaphase and the distance from the center. (D) Quantification of percentage of cell-division angle data. The angle is divided into three groups (0–30°, 30–60°, and 60–90°), and then the % of each group in different area sizes is shown (C). The angle of cell-division at different sizes during ectoderm differentiation (ECT) and undifferentiated (UND) cells is shown in the left and right columns, respectively.

To further analyze the timing, location, and direction of cell division, we used the histone H2B-EGFP knock-in iPSC line generated in our previous study.^[41]^ These cells were seeded in a pattern culture and maintained in an undifferentiated medium or neural differentiation medium. Time-lapse images were captured every 10 min for more than 16 h (**Figure 5B, Movie S1**). Taking advantage of the structure that is formed when the H2B chromosomes line up in the metaphase plane during cell division, the angle between the metaphase plate of the H2B-EGFP and the center of the pattern culture was measured (**Figure 5C**).^[42–45]^ The angle between the center of the metaphase plate and the center of the pattern culture was classified as 0–30°, 30– 60°, and 60–90°. Under the undifferentiated medium condition, 0–30°, 30–60°, and 60–90° were almost equally distributed in the 800 and 1600 µm diameter seeding patterns, while 0–30° increased in those of 200 and 400 µm (**Figure 5D**). Conversely, under conditions of neural differentiation, the proportion of H2B-EGFP at 60–90° increased significantly for all diameter patterns. These results indicate that during neuronal differentiation, the angle orientation of cell division is tuned in a cell-autonomous manner, dividing the cells into the inner and outer layers of the pattern.

### 2.5. Effects of teratogens on neuronal differentiation systems in patterned cell seeding systems

HiPSC-derived neuronal differentiation systems are useful safety tests for neurodevelopmental toxicity.^[46–48]^ Therefore, we aimed to demonstrate whether this micropatterned neuronal differentiation system can be used as a model to study neurogenic toxicity. In this study, valproic acid (VPA; sodium valproate or 2-propylpentanoate), a common antiepileptic drug with strong teratogenic properties, was used as a test substance.^[49–53]^ The PAX6-TEZ line was seeded on the micropattern culture using the LILAC-P system and differentiated into the neuroectoderm, as described above. During differentiation, the cells were treated with 0, 1, 3, or 10 mM VPA. Treatment with 10 mM VPA resulted in major cell death (**Figure 6A**). Therefore, the analysis was performed using 0, 1, and 3 mM VPA. Under these conditions, PAX6-tdTomato-positive cells were uniformly obtained after one week (**Figure 6B**). VPA had no effect on the induction of the differentiation of PAX6-positive neural progenitors. These cells were also examined for N-cadherin expression by immunocytochemistry (**Figure 6C**). The fluorescence intensity profiles for each pattern were plotted as a line and analyzed (**Figure 6D**). The fluorescence intensity at the circle micropatterns of 1600, 800, and 400 was enhanced in the 3 mM VPA treatment conditions (**Figure 6E**). Conversely, for the 200 µm diameter area, the fluorescence intensity decreased with VPA treatment. Thus, VPA affects the self-organization of micropatterned cultures of neuroectodermal cells, and its mode of action is dependent on the size of the micropatterned culture.

**Figure 6.**
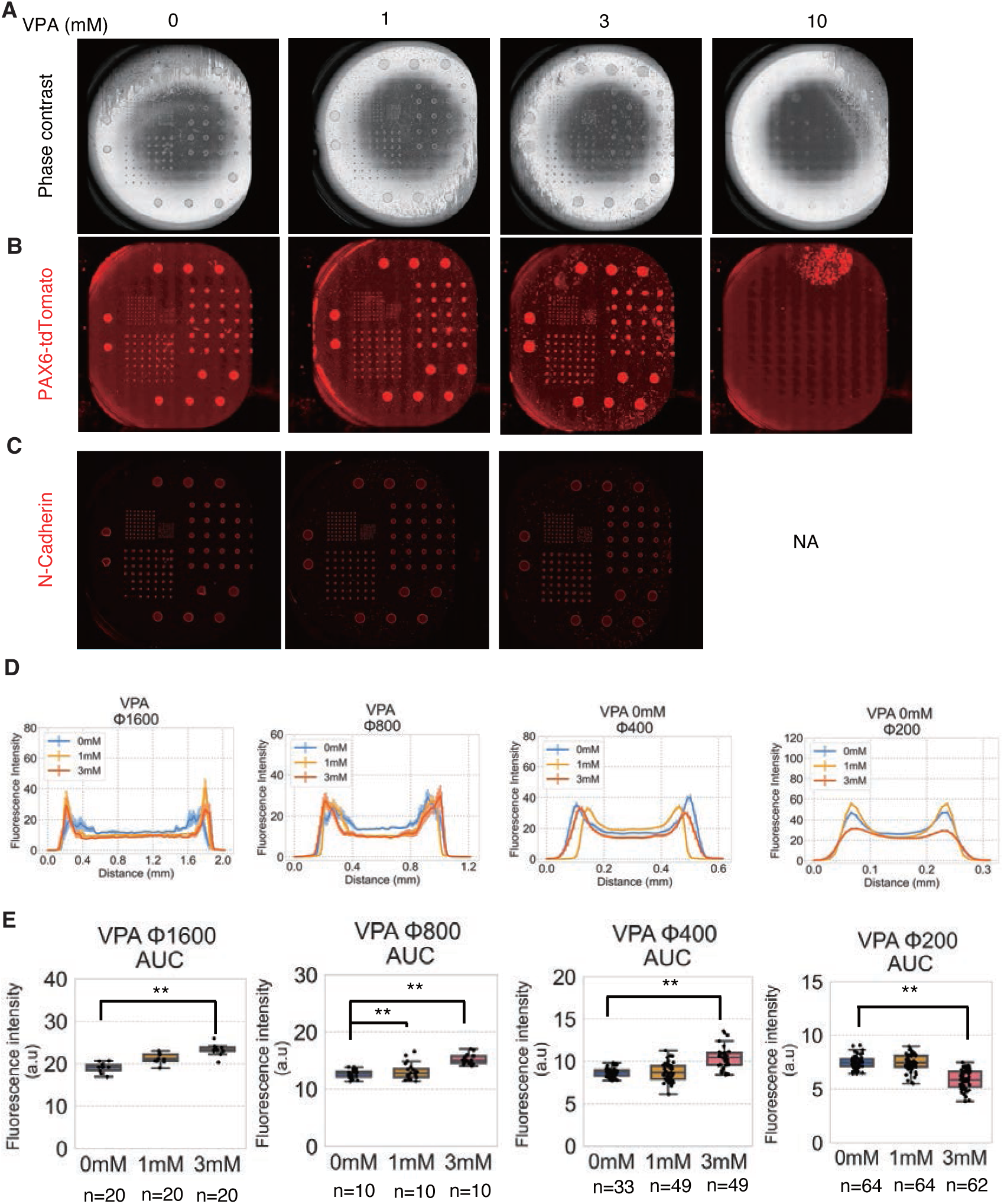
Effect of VPA on micropatterned neural structures formed with LILAC-P. (A) Representative whole dish images with phase contrast microscopes. hiPSCs were treated with 0, 1, 3, or 10 mM VPA for 6 days. (B) Representative whole dish images with PAX6-tdTomato fluorescence (red). (C) Representative whole dish images with N-cadherin expression detected using immunocytochemistry. (D) Averaged fluorescence intensity of N-cadherin expression in line distance of the micropatterned images. (E) Averaged fluorescence intensity of N-cadherin expression measured along the line distance of the micropatterned images. Bar graphs show the mean ± SE. ** indicate p<0.01 in the statistical test. P-values were determined by Dunnett’s test relative to non-treated conditions (0 mM).

## 3. Discussion

In this study, we developed a technology that automatically irradiated the entire culture dish following a designed image data input and succeeded in cell patterning corresponding to the customized design. This technology is based on the principle that ECM proteins are denatured only in the laser-irradiated area of culture dishes coated with a light-responsive polymer. LILAC-P does not require specialized knowledge and skills for the fabrication of micropatterning and allows flexible execution of new experiments using newly designed template for cell patterning. Furthermore, scalable patterning for mass cell culture was achievable using a high-speed, low-energy laser, making it applicable to regenerative medicine, which requires a large number of cells. Although in this study we focused on hiPSCs, our method can be applied in various biomedical fields, including basic research, drug development, and cell therapy, because it can be used, in principle, with various ECMs and cell types.

This method is enabled by thermally denaturing the ECM on a culture dish coated with a photoresponsive substrate using high-speed laser irradiation. Previous studies have reported methods wherein the surface properties of the culture substrate are modified by laser irradiation to alter the ECM deposition and the cell seeding pattern.^[54–59]^ Only one study directly heat-denatured the ECM using high-power lasers to ensure that its cell adhesion capacity was altered.^[60]^ However, these methods rely on high-energy lasers focused at localized areas (typically at the single-to hundred-cell levels), making it difficult to perform processing of entire cell cultures. In this study, we achieved highly efficient photothermal conversion using a photoresponsive substrate and precise processing over a large culture area with a short irradiation time using a high-speed laser of over 100 mm s^−1^ and less than 1 W power. Laser irradiation denatures recombinant laminin and causes it to lose its ability to bind to integrin, the major receptor on the cell surface; however, it does not inhibit subsequent cell or protein adhesion. Therefore, when cells secrete ECM and posess high migration ability, they deviate from the initial pattern during the course of culture. This suggests that it may be possible to analyze cell migration using this method. In addition, when ECM components, such as serum, are present in the culture medium after laser irradiation, caution is required because they may interfere with the formation of the seeding pattern.

The present study showed that reproducible cell-autonomous patterns of neuronal differentiation is achievable, especially neural differentiation, using circular seeding patterns of several diameters. Such patterns of cellular differentiation have been found in structures called neural rosettes and are also spontaneously seen during neural differentiation in two-dimensional cultures.^[61–65]^ However, their degree of formation and size are usually random, and the reproducibility of tissue formation patterns in each organoid is poor. Three-dimensional brain organoids are advanced models of brain development. However, the intracellular observation of their living cells is challenging. Only two studies have demonstrated that micropatterning systems can generate uniform neural rosettes using specialized micropatterned glass coverslips or slides.^[66–67]^ In this study, we showed that seeding cells in a circular pattern of several diameters in polystyrene dishes promotes neural differentiation and the spontaneous formation of a relatively reproducible neural layered structure. After neural induction for up to 1 week, PAX6 expression was induced more rapidly and uniformly than in simple monolayer cultures. In addition, ring-like structures with cells that were strongly positive for N-cadherin were generated. These findings are consistent with previous studies that used different micropatterning system.^[66–67]^ We further showed that an inward-to-outward orientation of nestin, a neuroprogenitor cell-specific intermediate diameter filament, depending on the pattern diameter. In addition, in cultures maintained up to 3 weeks, the expression of cortical neuronal markers TBR1, TBR2, TUJ1, BRN2, and Reelin was promoted from a relatively early stage, with TUJ1, SOX2, and BRN2 exhibiting ring-like expression localized to the outer region of the pattern. To explore the cellular basis of these self-organizing structures, we examined the localization and orientation of cell division in patterned cultures based on the behavior of pH3-expressing cells and H2B-EGFP in live cells. Such oriented and uneven localization of expressed molecules in micropattern cultures may result in biased cell signaling, asymmetric division, and generation of a self-organized layer structure during neural development.

We also demonstrated that our system can be applied as a model for neurodevelopmental toxicity. For example, we demonstrated that VPA treatment altered N-cadherin expression patterns depending on cell size. VPA has been an effective antiepileptic drug for many years; however, it is teratogenic and may be associated with neurodevelopmental delay and autism spectrum disorders in children of women exposed to the drug during pregnancy.^[68–70]^ Therefore, VPA is commonly used as a positive control for neurodevelopmental toxicity *in vitro*. VPA has pleiotropic effects on neuronal differentiation from hPSCs, depending on the cellular status and conditions, through epigenetic regulations.^[52–53, 71–77]^ Our findings indicate that the effect of VPA on the shape of N-cadherin expression changes with different micropattern sizes, which may reflect the pleiotropic effects of VPA. Further studies are warranted to examine the molecular mechanisms by which VPA affects cells cultured in micropatterns of different sizes and layers. In addition, a high-throughput evaluation of the effects of various substances would increase the usefulness of LILAC-P as a neurotoxicity evaluation test.

## 4. Conclusion

In this study, we developed an innovative micropatterning technique, LILAC-P, which is lithography-, hydrogel-, and PDMS-free. LILAC-P allows on-demand patterning by heat inactivation of the ECM on culture substrates using high-speed laser irradiation. This technology is expected to minimize the heterogeneity in the microenvironment of cultured cells. We believe that this micropatterned cell culture technique will contribute to elucidate the basic mechanisms of cell-autonomous tissue formation, investigate the effects of toxic substances, and refine tissue formation in regenerative medicine.

## 5. Experimental Section

### 5.1. Fabrication of photoresponsive culture substrates

The fabrication of photoresponsive culture substrates is described in our previous study.^[28]^ Briefly, the photoresponsive poly[(methyl methacrylate)-co-(Disperse Yellow 7 methacrylate)] (#579149, Sigma-Aldrich, St. Louis, MO, USA), which contains ∼25 mol% azobenzene moieties, was dissolved in a mixture of 2,2,2-trifluoroethanol and 1,1,1,3,3,3-hexafluoro-2-propanol (10 wt%) to make a 1.0 wt% solution. Next, 20 μL of the polymer solution was spin-coated on a 35 mm diameter polystyrene cell culture dish (#3000-035, AGC Techno Glass, Japan) at 2,000 rpm under an N_2_ atmosphere. The coated substrates were then annealed for 2 h at 80 °C. The absorbance of the substrate was typically 0.25 at 405 nm, and < 0.05 at wavelengths exceeding 500 nm.

### 5.2. Cell culture

HiPSCs (201B7, 454E2, and 1383D6 lines)^[32, 78–79]^ were obtained from CiRA (Center of iPS Cell Research and Application), Kyoto University, through the RIKEN Bioresource Research Center (Tsukuba, Japan). The WTC11 hiPSC line^[80–81]^ was obtained from Prof. Bruce Conklin at the Gladstone Institute of Cardiovascular Diseases, Coriell Institutes. H2B-GEZ (derived from the 1383D6 line) ^[41]^ and PAX6-TEZ (derived from the 454E2 line) ^[38]^ were generated in our laboratory as described in previous studies. These hiPSC lines were routinely cultured in StemFit AK02N medium (AK02N, Ajinomoto, Tokyo, Japan) ^[79]^ supplemented with its recommended supplements on 0.25 µg cm^−2^ recombinant laminin 511 fragments (iMatrix-511 silk, 892021, Nippi, Tokyo, Japan)-coated dishes^[82–86]^ A Rho-associated protein kinase (ROCK) inhibitor, Y-27632 (034-24024, Wako, Tokyo, Japan) (10 µM) was added to the passage medium.^[85, 87]^ For seeding hiPSCs onto the laser-irradiated cells, hiPSCs adhered to culture dishes were washed with PBS and then treated with TrypLE Select (A1285901, Thermo Fisher Scientific) at 37 °C for approximately 10 min. After aspirating the solution, the cells were resuspended in a fresh medium. hiPSCs were collected by cell scraping and pipetting. After counting the cell numbers, the cells were seeded at a density of 1 × 10^6^ cells per dish in LILAC-P-treated dishes. For differentiation into neuroectoderm, the medium was changed to StemFit AK02N medium without supplement C (StemFit w/o C) supplemented with 10 µM DMH1 (041-33881, Fujifilm Wako) and 10 µM SB431542 (192-16543, Fujifilm Wako) one day after seeding (day 1). The medium was changed every alternate day beginning on day 1 after seeding. On day 7, the medium was changed to neurobasal medium (21103049, Thermo Fisher Scientific) with 1x N-2 supplement (17502048, Thermo Fisher Scientific), 20 ng mL^−1^ EGF (058-09521, Fujifilm Wako), and 20 ng mL^−1^ FGF2 (064-05384, Fujifilm Wako) for further differentiation into neuronal lineages until day 21. For differentiation into mesoderm, the medium was changed to StemFit w/o C supplemented with 3 µM CHIR99021 (038-23101, Fujifilm Wako) and 10 ng mL^−1^ BMP4 (020-18851, Fujifilm Wako) one day after seeding (day 1). The medium was changed every alternate day beginning on day 1 after seeding. On day 5, the medium was replaced with StemPro-34 SFM (1X) (10639011, Thermo Fisher Scientific) supplemented with 100 ng mL^−1^ VEGF-A (225-02471, Fujifilm Wako) and cultured for 2 days. For differentiation into endoderm, the medium was changed to StemFit w/o C supplemented with 3 µM CHIR99021 and 10 ng mL^−1^ Activin A. On day 5, the medium was changed to StemSure D-MEM (197-16275, Fujifilm Wako) supplemented with 15% StemSure Serum Replacement (191-18375, Fujifilm Wako), 1% MEM non-essential amino acid solution (139-15651, Fujifilm Wako), 1% monothioglycerol solution (50 mM, x100) (195-15791, Fujifilm Wako), 1% L-alanyl-L-glutamine solution (200 mM, x100) (016-21841, Fujifilm Wako), and 1% dimethyl sulfoxide (DMSO; D2650, Sigma-Aldrich).

### 5.3. Laser irradiation

To achieve high-speed automatic cell processing using 405 nm laser scanning, we developed an all-in-one apparatus capable of capturing phase contrast and fluorescence images and performed tiled, whole-dish laser scanning in an on-demand manner.^[28]^ The laser beam was irradiated by a semiconductor laser source device through an optical fiber waveguide attached to the exit lens of the optical driving system. Laser scanning was performed by motor-driven regulation of the exit lens along a coordinate axis in the horizontal direction with high speed and accuracy. The laser was focused at the cell-substrate interface by the optical driving system to ensure precise photothermal conversion by the light-responsive polymer. A phase-contrast microscopy system was used, set with a ring slit and an objective lens carrying a phase plate. Microscopic images were acquired using a complementary metal oxide semiconductor (CMOS) camera. The ON/OFF ratio of laser irradiation was regulated by the input electricity of the semiconductor laser and adjusted to the laser scanning point. The position of the electricity switch was determined based on the coordinates of the phase-contrast microscope and laser device.

Laser irradiation was conducted at a specific energy [W] and speed [mm s^−1^] in a laser interval [pitch]. The design (**Fig. 1B**) made using Photoshop was saved in a TIFF format and then read into the machine software program. After pre-coating with iMatrix (0.5 µg cm^−2^), the dish was replaced with PBS and incubated at 37 °C for 1 h. Laser irradiation was performed using a CPD-017 cell-processing device (Kataoka Corporation). To determine the appropriate conditions for laser irradiation, the dishes were automatically irradiated as per the design by changing the laser power to 0.1, 0.2, 0.4, and 0.8 W. Laser irradiation was conducted twice in the same dish: first at 120 mm s^−1^ with a pitch interval of 20 µm, followed by a second irradiation under similar conditions but with a 30 µm pitch interval. To determine the appropriate laser irradiation speed, dishes were irradiated at speeds of 120, 240, and 480 mm s^−1^ at 0.8 W. The dish was irradiated twice as described above. For the detection of laminin and culture cells, laser irradiation was conducted at 0.8 W and 120 mm s^−1^, with a pitch interval of 20 µm at first, then irradiated under similar conditions but with a pitch interval of 30 µm. The irradiated culture dishes were then re-incubated and subjected to assays.

### 5.4. Solid-phase assays to detect the binding activity of laminin and fluorescence-labeled integrin

One tube (50 µg) of recombinant human integrin alpha 6 beta 1 protein (7809-A6, R&D Systems, USA) was labeled with HiLyte Fluor 647 Labeling Kit - NH2 (345-91051, Dojindo, Japan) according to the manufacturer’s protocol. Photoresponsive-substrate-coated culture dishes were pre-coated with iMatrix-511 silk (0.5 µg cm^−2^, Nippi, Japan) at 37 °C for 1 h. The dish was replaced with PBS(-) and then incubated at 4 °C for 1 h or more until the laser irradiation. After laser irradiation, dishes were washed twice with HBSS(+). Fluor 647-conjugated integrin (1:50, Integrin-647) was added to the irradiated dish and incubated at 25 °C for 2 h in the dark. The dishes were then washed twice with HBSS (+). Integrin expression was detected using a fluorescence microscope (BZ-X800; Keyence). Fluorescence intensity was calculated using ImageJ software (https://imagej.net/ij/). Integrin-647 added to a non-irradiated dish served as a positive control, and non-labeled integrin added to an irradiated dish servedas a negative control.

### 5.5. Solid-phase assays to detect biotinylated laminin with fluorescent streptavidin

In total, 100 µg of the recombinant laminin (iMatrix-511 silk) was biotinylated with Biotin Labeling Kit - NH2 (344-91141, Dojindo) according to the manufacturer’s protocol. Photoresponsive-substrate-coated culture dishes were pre-coated with 1 µg cm^−2^ biotinylated laminin at 37 °C for 1 h. The dishes were replaced with PBS(-) and then incubated at 4 °C for 1 h or more until the laser irradiation. After laser irradiation, dishes were washed twice with HBSS (+). Alexa Fluor 488 streptavidin (1:50, 405235, Biolegend, USA) was then added to the irradiated dish and incubated at room temperature for 2 h in the dark. The dishes were then washed twice with HBSS (+). The fluorescent streptavidin-biotin-laminin binding was detected using a fluorescence microscope (BZ-X800; Keyence). Fluorescence intensity was calculated using ImageJ software. Biotinylated laminin and fluorescent streptavidin were added to non-irradiated dishes as positive controls. The addition of non-biotinylated laminin and fluorescent streptavidin to the irradiated dish served as a negative control.

### 5.6. Immunocytochemistry

The expression of self-renewal and differentiation markers was detected by immunocytochemistry as previously described.^[88–94]^ Cells were fixed in PBS containing 4% (v v^−1^) paraformaldehyde (09154-85, Nacalai Tesque) for 10 min at room temperature. Cells were permeabilized with PBS containing 0.1% Triton X-100 (35501-02, Nacalai Tesque) for 10 min at room temperature, washed with PBS, and blocked with 0.1% BSA or 0.1% FBS. Subsequently, the cells were incubated with primary antibodies (listed in Table S1) at 4 °C overnight. After washing with PBS four times, the cells were incubated with secondary antibodies (listed in Table S2) at room temperature for 1 h or 4 °C overnight. After washing with PBS three times, cell nuclei were stained with the 4’,6-diamidino-2-phenylindole (DAPI) contained in the Fluoro-KEEPER antifade reagent, non-hardening type with DAPI (12745-74, Nacalai Tesque). Randomly selected images were analysed using a BZ-X800 microscope (Keyence, Osaka, Japan). Fluorescence intensity was calculated using ImageJ software.

## Data availability

All data generated or analyzed during this study are included in this published article (and its Supplementary Information files).

## Author Contributions

Y.H., K.S., and J.M. initiated the study. Y.H. developed the hiPSC assays and analyzed the data. T.W., K.S., and J.M. developed the fabrication process for the photoresponsive polymer substrates. J.M. developed the laser irradiation, image acquisition, and analysis system, and M.S., Y.H., K.H., and K.S. supervised the study and wrote the manuscript. Y.H. drafted the manuscript. All the authors have edited and approved the manuscript.

## Acknowledgements

We thank Dr. Bruce Conklin for providing us with the WTC11 hiPSC line, and Dr. Shinya Yamanaka, Dr. Masato Nakagawa, and Dr. Keisuke Okita for providing us with the 201B7, 454E2, and 1383D6 hiPSC lines. We express our sincere gratitude to Dr. Tamami Wakabayashi, Dr. Mami Matsuo-Takasaki, Ms. Yasuko Hemmi, and Ms. Iori Sato at RIKEN BRC for their technical assistance.

Received: ((will be filled in by the editorial staff))

Revised: ((will be filled in by the editorial staff))

Published online: ((will be filled in by the editorial staff))

## Supporting Information

Supporting Information is available from the Wiley Online Library or from the author.

## References

[1] F. Klemm, J. A. Joyce, Trends Cell Biol 2015, 25, 198 10.1016/j.tcb.2014.11.006.

[2] W. Sun, J. Lv, S. Guo, M. Lv, Front Cell Dev Biol 2023, 11, 1323678 10.3389/fcell.2023.1323678.

[3] M. Abdul-Al, G. K. Kyeremeh, M. Saeinasab, S. Heidari Keshel, F. Sefat, Bioengineering (Basel) 2021, 8 10.3390/bioengineering8080108.

[4] K. Kise, Y. Kinugasa-Katayama, N. Takakura, Adv Drug Deliv Rev 2016, 99, 197 10.1016/j.addr.2015.08.005.

[5] E. W. Young, D. J. Beebe, Chem Soc Rev 2010, 39, 1036 10.1039/b909900j.

[6] Z. Gu, J. Guo, H. Wang, Y. Wen, Q. Gu, Cell Prolif 2020, 53, e12754 10.1111/cpr.12754.

[7] T. Chavez, S. Gerecht, Trends Mol Med 2023, 29, 35 10.1016/j.molmed.2022.10.005.

[8] J. Barthes, H. Ozcelik, M. Hindie, A. Ndreu-Halili, A. Hasan, N. E. Vrana, Biomed Res Int 2014, 2014, 921905 10.1155/2014/921905.

[9] D. Park, J. Lim, J. Y. Park, S. H. Lee, Stem Cells Transl Med 2015, 4, 1352 10.5966/sctm.2015-0095.

[10] M. Thery, J Cell Sci 2010, 123, 4201 10.1242/jcs.075150.

[11] P. Srivastava, K. A. Kilian, Front Bioeng Biotechnol 2019, 7, 357 10.3389/fbioe.2019.00357.

[12] G. Blin, Development 2021, 148 10.1242/dev.186387.

[13] Y. Li, W. Jiang, X. Zhou, Y. Long, Y. Sun, Y. Zeng, X. Yao, Yale J Biol Med 2023, 96, 527 10.59249/UXOH1740.

[14] D. Falconnet, G. Csucs, H. M. Grandin, M. Textor, Biomaterials 2006, 27, 3044 10.1016/j.biomaterials.2005.12.024.

[15] G. Chen, N. Kawazoe, Adv Exp Med Biol 2020, 1250, 141 10.1007/978-981-15-3262-7_10.

[16] J. J. Norman, T. A. Desai, Ann Biomed Eng 2006, 34, 89 10.1007/s10439-005-9005-4.

[17] J. Christoffersson, C. F. Mandenius, Methods Mol Biol 2019, 1994, 227 10.1007/978-1-4939-9477-9_21.

[18] H. Otsuka, Molecules 2010, 15, 5525 10.3390/molecules15085525.

[19] T. H. Park, M. L. Shuler, Biotechnol Prog 2003, 19, 243 10.1021/bp020143k.

[20] K. Wilson, M. Stancescu, M. Das, J. Rumsey, J. Hickman, J Vac Sci Technol B Nanotechnol Microelectron 2011, 29, 21020 10.1116/1.3549127.

[21] J. Y. Lee, S. S. Shah, C. C. Zimmer, G. Y. Liu, A. Revzin, Langmuir 2008, 24, 2232 10.1021/la702883d.

[22] J. L. Curley, S. R. Jennings, M. J. Moore, J Vis Exp 2011, 10.3791/2636.

[23] L. M. Murray, V. Nock, J. J. Evans, M. M. Alkaisi, J Nanobiotechnology 2014, 12, 60 10.1186/s12951-014-0060-6.

[24] W. Zheng, W. Zhang, X. Jiang, Adv Healthc Mater 2013, 2, 95 10.1002/adhm.201200104.

[25] H. Shin, Biomaterials 2007, 28, 126 10.1016/j.biomaterials.2006.08.007.

[26] H. J. Oh, J. Kim, H. Kim, N. Choi, S. Chung, Adv Healthc Mater 2021, 10, e2002122 10.1002/adhm.202002122.

[27] K. Hattori, S. Sugiura, T. Kanamori, Methods Mol Biol 2014, 1104, 251 10.1007/978-1-62703-733-4_17.

[28] Y. Hayashi, J. Matsumoto, S. Kumagai, K. Morishita, L. Xiang, Y. Kobori, S. Hori, M. Suzuki, T. Kanamori, K. Hotta, K. Sumaru, Commun Biol 2018, 1, 218 10.1038/s42003-018-0222-4.

[29] F. P. Nicoletta, D. Cupelli, P. Formoso, G. De Filpo, V. Colella, A. Gugliuzza, Membranes (Basel) 2012, 2, 134 10.3390/membranes2010134.

[30] P. Saraithong, P. Krajcarski, Y. Kusaka, M. Yamada, J. Matsumoto, H. Cunningham, S. Salih, D. Jones, D. Baddhan, C. Hausner, J. Anumonwo, A. Rosenzweig, M. M. Navarro, L. V. Diaz, J. Criscione, D. H. Kim, T. J. Herron, Commun Biol 2025, 8, 745 10.1038/s42003-025-08162-0.

[31] P. A. Mould, Methods Mol Biol 2009, 522, 195 10.1007/978-1-59745-413-1_13.

[32] K. Takahashi, K. Tanabe, M. Ohnuki, M. Narita, T. Ichisaka, K. Tomoda, S. Yamanaka, Cell 2007, 131, 861 10.1016/j.cell.2007.11.019.

[33] J. Dobner, S. Diecke, J. Krutmann, A. Prigione, A. Rossi, Nat Commun 2024, 15, 8547 10.1038/s41467-024-52922-1.

[34] S. M. Chambers, C. A. Fasano, E. P. Papapetrou, M. Tomishima, M. Sadelain, L. Studer, Nat Biotechnol 2009, 27, 275 10.1038/nbt.1529.

[35] C. L. Curchoe, J. Russo, A. V. Terskikh, Stem Cell Res 2012, 8, 239 10.1016/j.scr.2011.11.003.

[36] E. Frullanti, S. Amabile, M. G. Lolli, A. Bartolini, G. Livide, E. Landucci, F. Mari, F. M. Vaccarino, F. Ariani, L. Massimino, A. Renieri, I. Meloni, Eur J Hum Genet 2016, 24, 252 10.1038/ejhg.2015.79.

[37] J. Mariani, G. Coppola, P. Zhang, A. Abyzov, L. Provini, L. Tomasini, M. Amenduni, A. Szekely, D. Palejev, M. Wilson, M. Gerstein, E. L. Grigorenko, K. Chawarska, K. A. Pelphrey, J. R. Howe, F. M. Vaccarino, Cell 2015, 162, 375 10.1016/j.cell.2015.06.034.

[38] M. Matsuo-Takasaki, S. Kambayashi, Y. Hemmi, T. Wakabayashi, T. Shimizu, Y. An, H. Ito, K. Takeuchi, M. Ibuki, T. Kawashima, R. Masayasu, M. Suzuki, Y. Kawai, M. Umekage, T. M. Kato, M. Noguchi, K. Nakade, Y. Nakamura, T. Nakaishi, N. Nishishita, M. Tsukahara, Y. Hayashi, Elife 2024, 12 10.7554/eLife.89724.

[39] T. C. Schulz, G. M. Palmarini, S. A. Noggle, D. A. Weiler, M. M. Mitalipova, B. G. Condie, BMC Neurosci 2003, 4, 27 10.1186/1471-2202-4-27.

[40] R. J. McEvilly, M. O. de Diaz, M. D. Schonemann, F. Hooshmand, M. G. Rosenfeld, Science 2002, 295, 1528 10.1126/science.1067132.

[41] K. Nakade, S. Tsukamoto, K. Nakashima, Y. An, I. Sato, J. Li, Y. Shimoda, Y. Hemmi, Y. Miwa, Y. Hayashi, Cell Rep Methods 2023, 3, 100662 10.1016/j.crmeth.2023.100662.

[42] P. Meraldi, V. M. Draviam, P. K. Sorger, Dev Cell 2004, 7, 45 10.1016/j.devcel.2004.06.006.

[43] C. H. Tan, I. Gasic, S. P. Huber-Reggi, D. Dudka, M. Barisic, H. Maiato, P. Meraldi, Elife 2015, 4 10.7554/eLife.05124.

[44] T. Kanda, K. F. Sullivan, G. M. Wahl, Curr Biol 1998, 8, 377 10.1016/s0960-9822(98)70156-3.

[45] M. Charnley, F. Anderegg, R. Holtackers, M. Textor, P. Meraldi, PLoS One 2013, 8, e66918 10.1371/journal.pone.0066918.

[46] T. C. Stummann, L. Hareng, S. Bremer, Toxicology 2009, 257, 117 10.1016/j.tox.2008.12.018.

[47] H. T. Hogberg, J. Bressler, K. M. Christian, G. Harris, G. Makri, C. O’Driscoll, D. Pamies, L. Smirnova, Z. Wen, T. Hartung, Stem Cell Res Ther 2013, 4 Suppl 1, S4 10.1186/scrt365.

[48] A. Sachinidis, W. Albrecht, P. Nell, A. Cherianidou, N. J. Hewitt, K. Edlund, J. G. Hengstler, Trends Mol Med 2019, 25, 470 10.1016/j.molmed.2019.04.003.

[49] J. Kim, S. H. Park, W. Sun, Int J Stem Cells 2024, 10.15283/ijsc24066.

[50] G. Pietrogrande, M. R. Shaker, S. J. Stednitz, F. Soheilmoghaddam, J. Aguado, S. D. Morrison, S. Zambrano, T. Tabassum, I. Javed, J. Cooper-White, T. P. Davis, T. J. O’Brien, E. K. Scott, E. J. Wolvetang, Mol Psychiatry 2024, 10.1038/s41380-024-02732-0.

[51] K. Lauschke, A. K. Rosenmai, I. Meiser, J. C. Neubauer, K. Schmidt, M. A. Rasmussen, B. Holst, C. Taxvig, J. K. Emneus, A. M. Vinggaard, Arch Toxicol 2020, 94, 3831 10.1007/s00204-020-02856-6.

[52] K. Cui, Y. Wang, Y. Zhu, T. Tao, F. Yin, Y. Guo, H. Liu, F. Li, P. Wang, Y. Chen, J. Qin, Microsyst Nanoeng 2020, 6, 49 10.1038/s41378-020-0165-z.

[53] N. Dreser, K. Madjar, A. K. Holzer, M. Kapitza, C. Scholz, P. Kranaster, S. Gutbier, S. Klima, D. Kolb, C. Dietz, T. Trefzer, J. Meisig, C. van Thriel, M. Henry, M. R. Berthold, N. Bluthgen, A. Sachinidis, J. Rahnenfuhrer, J. G. Hengstler, T. Waldmann, M. Leist, Arch Toxicol 2020, 94, 151 10.1007/s00204-019-02612-5.

[54] C. B. Herbert, T. L. McLernon, C. L. Hypolite, D. N. Adams, L. Pikus, C. C. Huang, G. B. Fields, P. C. Letourneau, M. D. Distefano, W. S. Hu, Chem Biol 1997, 4, 731 10.1016/s1074-5521(97)90311-2.

[55] Y. Nahmias, D. J. Odde, Nat Protoc 2006, 1, 2288 10.1038/nprot.2006.386.

[56] Y. L. Khung, S. D. Graney, N. H. Voelcker, Biotechnol Prog 2006, 22, 1388 10.1021/bp060115s.

[57] S. Iwanaga, Y. Akiyama, A. Kikuchi, M. Yamato, K. Sakai, T. Okano, Biomaterials 2005, 26, 5395 10.1016/j.biomaterials.2005.01.021.

[58] M. Yamato, C. Konno, S. Koike, Y. Isoi, T. Shimizu, A. Kikuchi, K. Makino, T. Okano, J Biomed Mater Res A 2003, 67, 1065 10.1002/jbm.a.10078.

[59] C. Melero, A. Kolmogorova, P. Atherton, B. Derby, A. Reid, K. Jansen, C. Ballestrem, J Vis Exp 2019, 10.3791/60092.

[60] W. F. Heinz, M. Hoh, J. H. Hoh, Lab Chip 2011, 11, 3336 10.1039/c1lc20204a.

[61] P. G. Wilson, S. S. Stice, Stem Cell Rev 2006, 2, 67 10.1007/s12015-006-0011-1.

[62] M. Miotto, M. Rosito, M. Paoluzzi, V. de Turris, V. Folli, M. Leonetti, G. Ruocco, A. Rosa, G. Gosti, Front Cell Dev Biol 2023, 11, 1134091 10.3389/fcell.2023.1134091.

[63] Y. Elkabetz, G. Panagiotakos, G. Al Shamy, N. D. Socci, V. Tabar, L. Studer, Genes Dev 2008, 22, 152 10.1101/gad.1616208.

[64] I. Kelava, M. A. Lancaster, Cell Stem Cell 2016, 18, 736 10.1016/j.stem.2016.05.022.

[65] A. Deglincerti, F. Etoc, M. C. Guerra, I. Martyn, J. Metzger, A. Ruzo, M. Simunovic, A. Yoney, A. H. Brivanlou, E. Siggia, A. Warmflash, Nat Protoc 2016, 11, 2223 10.1038/nprot.2016.131.

[66] T. Haremaki, J. J. Metzger, T. Rito, M. Z. Ozair, F. Etoc, A. H. Brivanlou, Nat Biotechnol 2019, 37, 1198 10.1038/s41587-019-0237-5.

[67] G. T. Knight, B. F. Lundin, N. Iyer, L. M. Ashton, W. A. Sethares, R. M. Willett, R. S. Ashton, Elife 2018, 7 10.7554/eLife.37549.

[68] S. Duncan, Curr Opin Neurol 2007, 20, 175 10.1097/WCO.0b013e32805866fb.

[69] R. Bromley, Reprod Toxicol 2016, 64, 203 10.1016/j.reprotox.2016.06.007.

[70] D. Zarate-Lopez, A. L. Torres-Chavez, A. Y. Galvez-Contreras, O. Gonzalez-Perez, Curr Neuropharmacol 2024, 22, 260 10.2174/1570159X22666231003121513.

[71] K. Meganathan, S. Jagtap, S. P. Srinivasan, V. Wagh, J. Hescheler, J. Hengstler, M. Leist, A. Sachinidis, Cell Death Dis 2015, 6, e1756 10.1038/cddis.2015.121.

[72] C. C. Miranda, T. G. Fernandes, S. N. Pinto, M. Prieto, M. M. Diogo, J. M. S. Cabral, Toxicol Lett 2018, 294, 51 10.1016/j.toxlet.2018.05.018.

[73] V. B. R. Konala, S. Nandakumar, H. Surendran, S. Datar, R. Bhonde, R. Pal, Toxicol Appl Pharmacol 2021, 433, 115792 10.1016/j.taap.2021.115792.

[74] V. C. de Leeuw, C. T. M. van Oostrom, P. F. K. Wackers, J. L. A. Pennings, H. M. Hodemaekers, A. H. Piersma, E. V. S. Hessel, Chemosphere 2022, 304, 13529810.1016/j.chemosphere.2022.135298.

[75] A. Chen, M. Wang, C. Xu, Y. Zhao, P. Xian, Y. Li, W. Zheng, X. Yi, S. Wu, Y. Wang, Front Mol Neurosci 2023, 16, 1151162 10.3389/fnmol.2023.1151162.

[76] F. Pistollato, D. Carpi, E. Mendoza-de Gyves, A. Paini, S. K. Bopp, A. Worth, A. Bal-Price, Reprod Toxicol 2021, 105, 101 10.1016/j.reprotox.2021.08.007.

[77] S. H. Schulpen, J. L. Pennings, A. H. Piersma, Toxicol Sci 2015, 146, 311 10.1093/toxsci/kfv094.

[78] K. Okita, Y. Matsumura, Y. Sato, A. Okada, A. Morizane, S. Okamoto, H. Hong, M. Nakagawa, K. Tanabe, K. Tezuka, T. Shibata, T. Kunisada, M. Takahashi, J. Takahashi, H. Saji, S. Yamanaka, Nat Methods 2011, 8, 409 10.1038/nmeth.1591.

[79] M. Nakagawa, Y. Taniguchi, S. Senda, N. Takizawa, T. Ichisaka, K. Asano, A. Morizane, D. Doi, J. Takahashi, M. Nishizawa, Y. Yoshida, T. Toyoda, K. Osafune, K. Sekiguchi, S. Yamanaka, Sci Rep 2014, 4, 3594 10.1038/srep03594.

[80] F. R. Kreitzer, N. Salomonis, A. Sheehan, M. Huang, J. S. Park, M. J. Spindler, P. Lizarraga, W. A. Weiss, P. L. So, B. R. Conklin, Am J Stem Cells 2013, 2, 119

[81] Y. Hayashi, E. C. Hsiao, S. Sami, M. Lancero, C. R. Schlieve, T. Nguyen, K. Yano, A. Nagahashi, M. Ikeya, Y. Matsumoto, K. Nishimura, A. Fukuda, K. Hisatake, K. Tomoda, I. Asaka, J. Toguchida, B. R. Conklin, S. Yamanaka, Proc Natl Acad Sci U S A 2016, 113, 13057 10.1073/pnas.1603668113.

[82] T. Miyazaki, S. Futaki, K. Hasegawa, M. Kawasaki, N. Sanzen, M. Hayashi, E. Kawase, K. Sekiguchi, N. Nakatsuji, H. Suemori, Biochem Biophys Res Commun 2008, 375, 27 10.1016/j.bbrc.2008.07.111.

[83] T. Miyazaki, S. Futaki, H. Suemori, Y. Taniguchi, M. Yamada, M. Kawasaki, M. Hayashi, H. Kumagai, N. Nakatsuji, K. Sekiguchi, E. Kawase, Nat Commun 2012, 3, 1236 10.1038/ncomms2231.

[84] T. Miyazaki, E. Kawase, Curr Protoc Stem Cell Biol 2015, 32, 1C 18 1 10.1002/9780470151808.sc01c18s32.

[85] T. Miyazaki, T. Isobe, N. Nakatsuji, H. Suemori, Sci Rep 2017, 7, 41165 10.1038/srep41165.

[86] E. Borisova, K. Nishimura, Y. An, M. Takami, J. Li, D. Song, M. Matsuo-Takasaki, D. Luijkx, S. Aizawa, A. Kuno, E. Sugihara, T. A. Sato, F. Yumoto, T. Terada, K. Hisatake, Y. Hayashi, iScience 2022, 25, 103525 10.1016/j.isci.2021.103525.

[87] M. Nakagawa, M. Koyanagi, K. Tanabe, K. Takahashi, T. Ichisaka, T. Aoi, K. Okita, Y. Mochiduki, N. Takizawa, S. Yamanaka, Nat Biotechnol 2008, 26, 101 10.1038/nbt1374.

[88] Y. Arai, H. Ito, T. Shimizu, Y. Shimoda, D. Song, M. Matsuo-Takasaki, T. Hayata, Y. Hayashi, Front Cell Dev Biol 2024, 12, 1370723 10.3389/fcell.2024.1370723.

[89] D. Takagi, S. Tsukamoto, K. Nakade, T. Shimizu, Y. Arai, M. Matsuo-Takasaki, M. Noguchi, Y. Nakamura, N. Yumoto, J. Kawada, T. Hayata, Y. Hayashi, Stem Cell Res 2024, 79, 103493 10.1016/j.scr.2024.103493.

[90] M. Mori, S. Yoshii, M. Noguchi, D. Takagi, T. Shimizu, H. Ito, M. Matsuo-Takasaki, Y. Nakamura, S. Takahashi, H. Hamada, K. Ohnuma, T. Shiohama, Y. Hayashi, Stem Cell Res 2024, 77, 103432 10.1016/j.scr.2024.103432.

[91] R. Li, H. Tsuboi, H. Ito, D. Takagi, Y. H. Chang, T. Shimizu, Y. Arai, M. Matsuo-Takasaki, M. Noguchi, Y. Nakamura, K. Ohnuma, S. Takahashi, Y. Hayashi, Stem Cell Res 2024, 81, 103584 10.1016/j.scr.2024.103584.

[92] N. Ge, K. Suzuki, I. Sato, M. Noguchi, Y. Nakamura, M. Matsuo-Takasaki, J. Fujishiro, Y. Hayashi, Hum Cell 2024, 38, 18 10.1007/s13577-024-01147-x.

[93] D. Song, G. Takahashi, Y. W. Zheng, M. Matsuo-Takasaki, J. Li, M. Takami, Y. An, Y. Hemmi, N. Miharada, T. Fujioka, M. Noguchi, T. Nakajima, M. K. Saito, Y. Nakamura, T. Oda, Y. Miyaoka, Y. Hayashi, Hum Mol Genet 2022, 31, 3652 10.1093/hmg/ddac080.

[94] T. Shimizu, M. Matsuo-Takasaki, D. Luijkx, M. Takami, Y. Arai, M. Noguchi, Y. Nakamura, T. Hayata, M. K. Saito, Y. Hayashi, Stem Cell Res 2022, 61, 102744 10.1016/j.scr.2022.102744.

