## Supplementary Information for "Micropatterned neural induction with heat-inactivated extracellular matrix protein by on-demand high-speed laser"

Table of content:
Supplementary Figures S1-5
Supplementary Tables S1-2

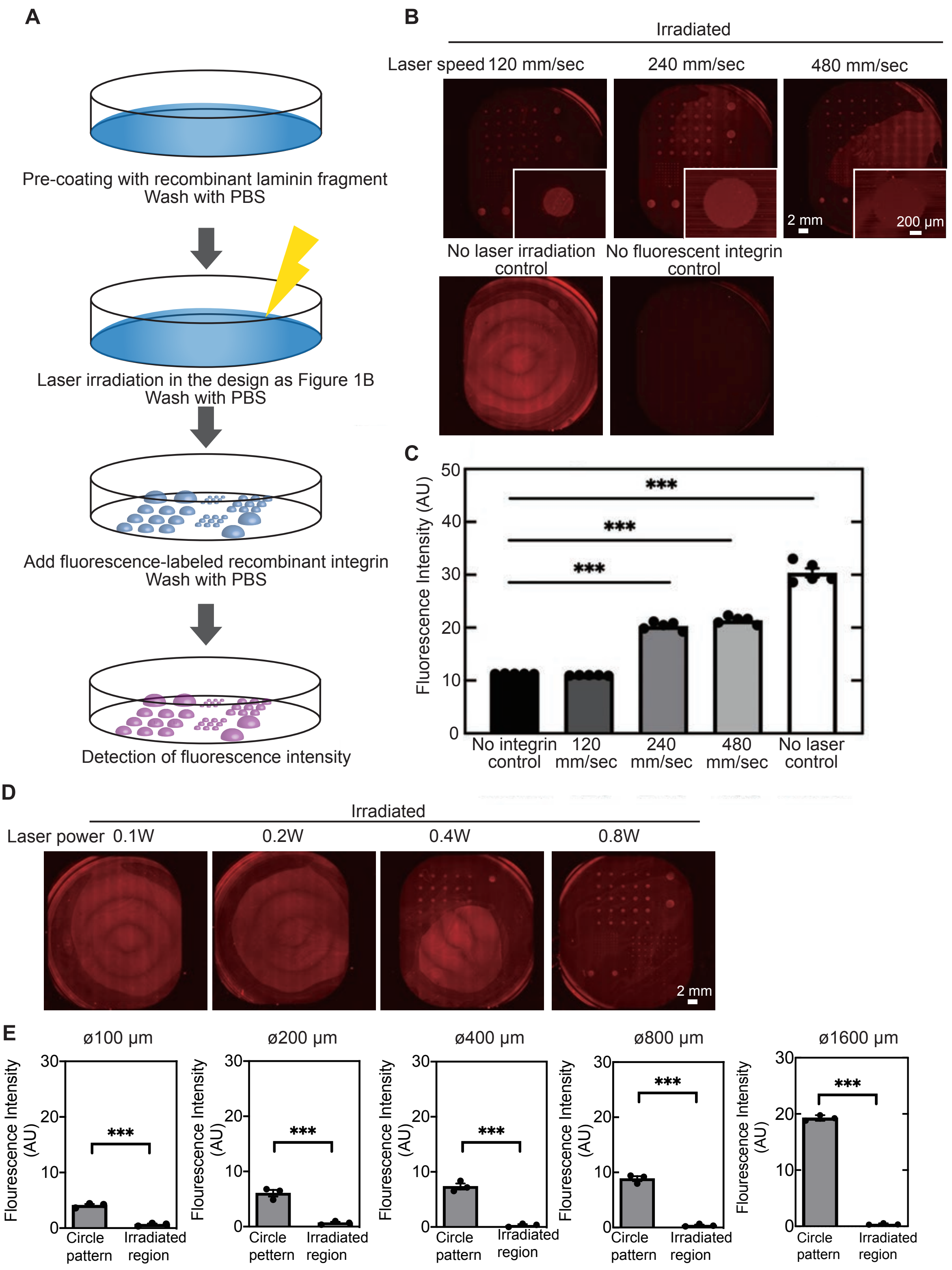


**Figure S1. Detection of laminin-integrin interaction inactivated with LILAC-P using solid-phase assays.** (A) The schematics of the detection of the extracellular matrix (ECM) proteins by fluorescent-labeled integrin alpha 6 beta 1. (B) Representative images of the laminin detection assay in the dishes with LILACP at laser speed of 120, 240, or 480 mm $S^{-1}$. Also, a positive control dish (no laser irradiation) and a negative control dish (no recombinant integrin treatment) are shown. (C) Quantification of fluorescent intensity of the laminin detection assay in the dish irradiating at 120, 240, or 480 mm $S^{-1}$. All data show the mean ± SEM (n=5). All groups are compared with the negative control condition. The p-values were determined by one-way ANOVA with Dunnett's multiple comparison test. *** indicate p<0.001 in the statistical test. (D) The examination of the appropriate laser power by changing at 0.1, 0.2, 0.4, and 0.8 W. Representative images were whole dishes in each laser power the laminin detection assay. (E) the quantification of the fluorescence intensity of integrin-647 in each size (⌀100, ⌀200, ⌀400, ⌀800, and ⌀1600μm). All data show the mean ± SEM (n=3). All groups are compared with the negative control condition. The p-values are determined by unpaired Student's t-test. *** indicate p<0.001 in the statistical test.

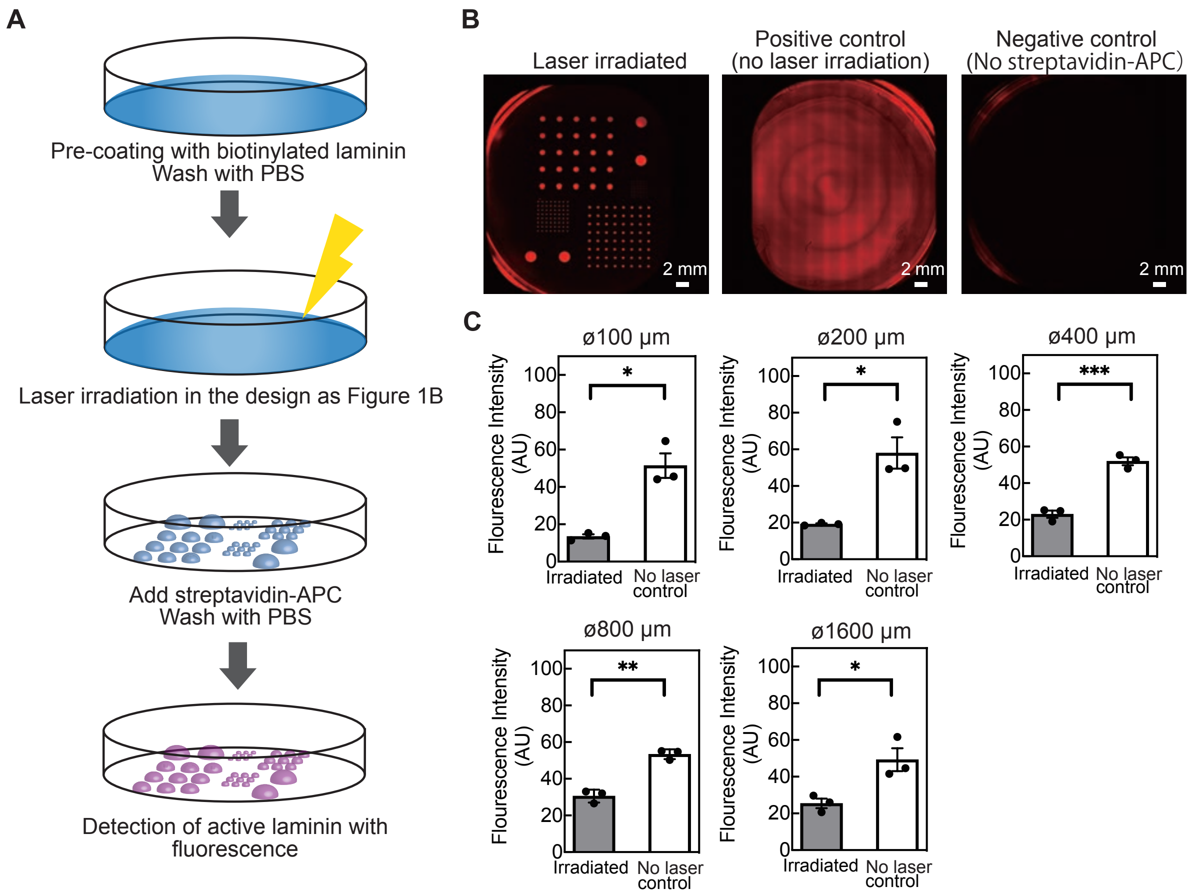


**Figure S2. Detection of denatured laminin with LILAC-P using solid-phase assays.** (A) The schematic process of the detection of the extracellular matrix (ECM) proteins by biotin-labeled laminin and APC-conjugated APC. (B) Representative images of the solid-phase assays. An optimized laser-irradiated dish (left image), a positive control (no laser-irradiated) dish (middle image), and a negative control (no streptavidin-APC treated) dish (right image) are shown. (C) Quantification of fluorescence intensity of the solid-phase assays. All data show the mean ± SEM (n=5). The p-values are determined by unpaired Student's t-test. *, **, and *** indicate p<0.05, p<0.01, and p<0.001 in the statistical tests, respectively.

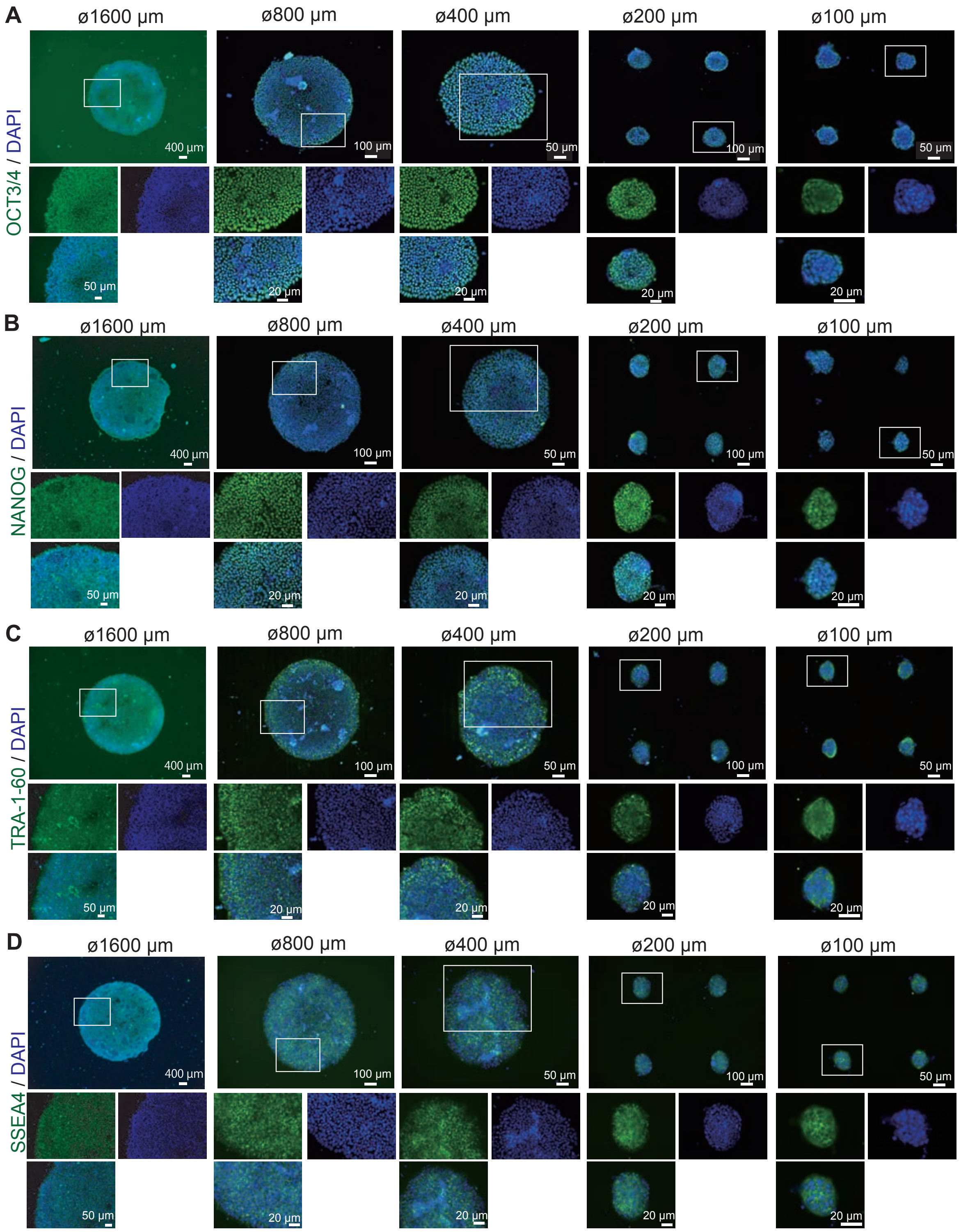


**Figure S3. Undifferentiated hiPSCs cultured on dishes micropatterned with LILAC-P.** HiPSCs cultured on LILACP-treated dishes for 2 days were examined with their self-renewal markers. Representative merged images show the expression of OCT 3/4 (A), NANOG (B), TRA-1-60 (C), and SSEA4 (D) (green) and DAPI (blue) at different sizes ⌀1600 µm, ⌀800 µm, ⌀400 µm, ⌀200 µm, and ⌀100 µm from left to right. Bottom panels, enlargements of inserts (white square) in upper panels and expression of each individual marker.

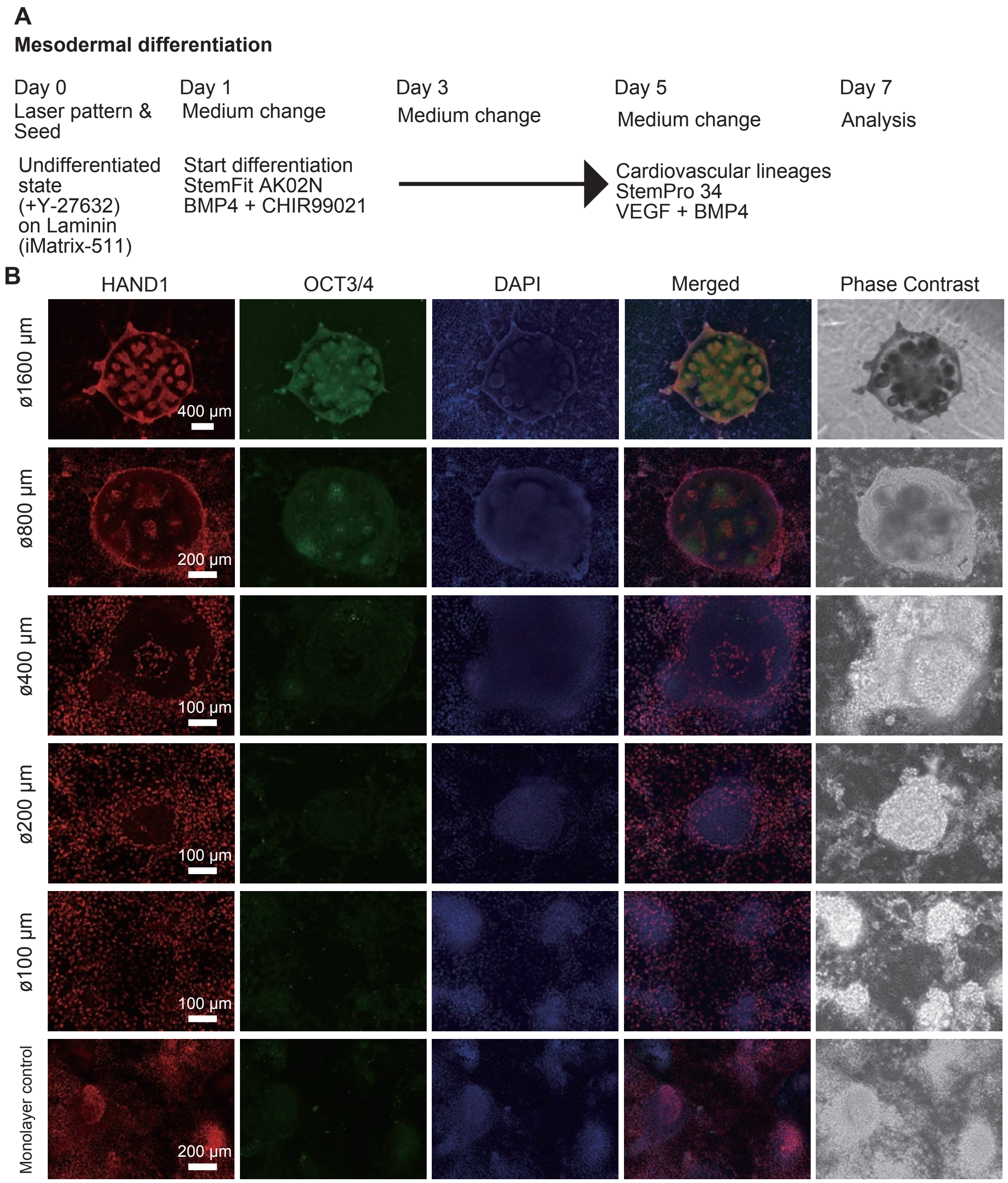


**Figure S4. Mesodermal differentiation from hiPSCs cultured on dishes micropatterned with LILAC-P.** (A) Schematics of mesodernal differentiation in culture dishes with LILACP system for one week. (B) Representative images of HAND1 (Red), OCT3/4 (Green), and DAPI (Blue) detected with immunocytochemistry. Merged and Phase contrast images from the right at different sizes (i.e., ⌀1600 µm, ⌀800 µm, ⌀400 µm, ⌀200 µm, ⌀100 µm) and monolayer control samples.

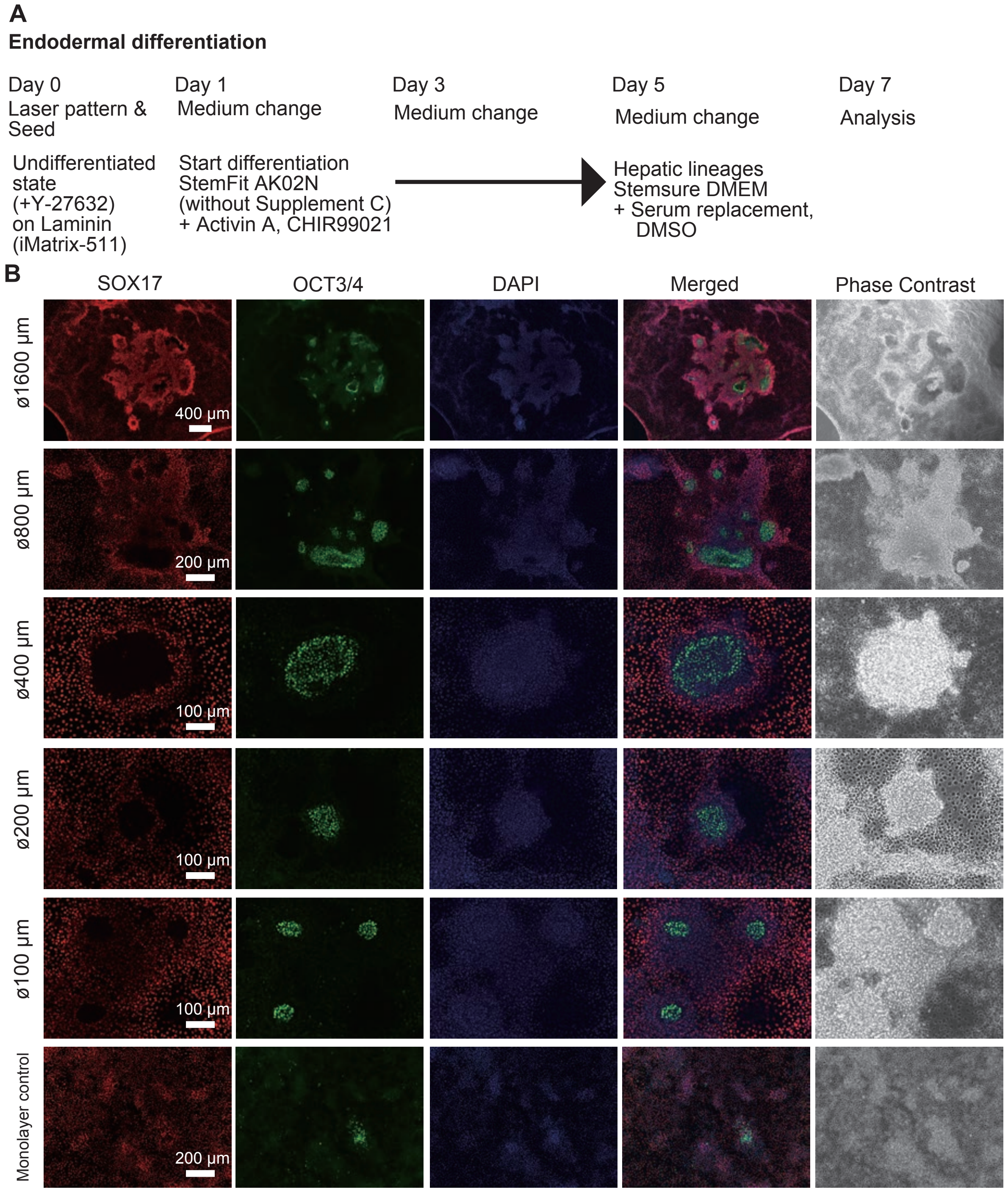


**Figure S5. Endodermal differentiation from hiPSCs cultured on dishes micropatterned with LILAC-P.** (A) Schematics of mesodernal differentiation in culture dishes with LILACP system for one week. (B) Representative images of HAND1 (Red), OCT3/4 (Green), and DAPI (Blue) detected with immunocytochemistry. Merged and Phase contrast images from the right at different sizes (i.e., ⌀1600 μm, ⌀800 μm, ⌀400 μm, ⌀200 μm, ⌀100 μm) and monolayer control samples.

**Table S1. The list of primary antibodies used in this study.**

| Name | Host species | Dilutions or Concentrations in immunocytochemistry | SOURCE | IDENTIFIER |
|---|---|---|---|---|
| NANOG | Rabbit | 0.5 μg/mL | Reprocell | Cat#RCAB004P-F |
| OCT-3/4 | Mouse | 0.4 μg/mL | Santa Cruz Biotechnology | Cat#sc-5279 |
| PAX6 | Rabbit | 1:400 | MBL | Cat#PD022 |
| SOX17 | Goat | 0.4 μg/mL | R&D systems | Cat#AF1924 |
| FITC Mouse anti-SSEA-4 | Mouse | 1:5 | BD bioscience | Cat#560126 |
| Alexa Fluor 488 anti-TRA-1-60 | Mouse | 1:10 | BD bioscience | Cat#560173 |
| TUJ1 | Mouse | 1 μg/mL | R&D systems | Cat#MAB1195 |
| FOXG1 | Rabbit | 1:200 | Abcam | Cat#ab18259 |
| N-Cadherin | Mouse | 2 μg/mL | Biolegend | Cat#350802 |
| Nestin | Mouse | 2 μg/mL | Biolegend | Cat#656802 |
| TBR1 | Rabbit | 1:100 | Abcam | Cat#ab31940 |
| TBR2 | Rabbit | 1:500 | Abcam | Cat#ab23345 |
| SOX2 | Mouse | 10 μg/mL | R&D systems | Cat#2018 |
| BRN2 | Rabbit | 1:1,000 | Cell Signaling Technology | Cat#12137 |
| Reelin | Mouse | 10 μg/mL | MBL | Cat#223-3 |
| Phospho-Histone H3 (Ser10) (6G3) | Mouse | 1:400 | Cell Signaling Technology | Cat#9706S |
| HAND1 | Goat | 5 μg/mL | R&D systems | Cat#AF368 |

**Table S2. The list of secondary antibodies used in this study.**

| Name | Host species | Dilutions | SOURCE | IDENTIFIER |
|---|---|---|---|---|
| Anti-Mouse IgG (H+L) Highly Cross-Adsorbed Secondary Antibody, Alexa Fluor Plus 488 | Donkey | 1:1,000 | Thermo Fisher Scientific | Cat#A-21202 |
| Anti-Mouse IgG (H+L) Highly Cross-Adsorbed Secondary Antibody, Alexa Fluor Plus 555 | Donkey | 1:1,000 | Thermo Fisher Scientific | Cat#A-31570 |
| Anti-Mouse IgG (H+L) Highly Cross-Adsorbed Secondary Antibody, Alexa Fluor 647 | Donkey | 1:1,000 | Thermo Fisher Scientific | Cat#A-31571 |
| DyLight 488 anti-rabbit IgG (minimal x-reactivity) | Donkey | 1:1,000 | BioLegend | Cat#406404 |
| Anti-Rabbit IgG (H+L) Cross-Adsorbed Secondary Antibody, Alexa Fluor 555 | Goat | 1:1,000 | Thermo Fisher Scientific | Cat#A21428 |
| DyLight 649 anti-rabbit IgG | Donkey | 1:1,000 | BioLegend | Cat#406406 |
| Anti-Goat IgG (H+L) Highly Cross-Adsorbed Secondary Antibody, Alexa Fluor Plus 488 | Donkey | 1:1,000 | Thermo Fisher Scientific | Cat#A32814 |
| Anti-Goat IgG (H+L) Highly Cross-Adsorbed Secondary Antibody, Alexa Fluor Plus 555 | Donkey | 1:1,000 | Thermo Fisher Scientific | Cat#A32816 |
| Anti-Goat IgG (H+L) Highly Cross-Adsorbed Secondary Antibody, Alexa Fluor Plus 647 | Donkey | 1:1,000 | Thermo Fisher Scientific | Cat#A32849 |
